# Distinct ATRX functions cooperate with 9-1-1 and CST complexes to safeguard replication and telomere integrity

**DOI:** 10.1101/2025.09.22.677761

**Authors:** Sandra Segura-Bayona, Marija Maric, Tohru Takaki, Zornitsa Manova, Shudong Li, Tyler H Stanage, Aurora I Idilli, Graeme Hewitt, Phil Ruis, Rhona Millar, Harshil Patel, Steven Howell, Panagiotis Kotsantis, Michael Howell, Simon J Boulton

**Affiliations:** DSB Repair Metabolism Laboratory, The Francis Crick Institute, 1 Midland Road, London NW1 1AT, UK; Genome Stability Group, King’s College London, School of Cancer & Pharmaceutical Sciences, Comprehensive Cancer Centre, London SE1 9RT, UK; Bioinformatics and Biostatistics STP, The Francis Crick Institute, 1 Midland Road, London NW1 1AT, UK; Proteomics STP, The Francis Crick Institute, 1 Midland Road, London NW1 1AT, UK; High Throughput Screening STP, The Francis Crick Institute, 1 Midland Road, London NW1 1AT, UK; Biomedical and Life Sciences, Faculty of Health and Medicine, Lancaster University, Lancaster LA1 4YQ, UK

**Keywords:** ATRX, CST complex, 9-1-1 complex, ATR, FAM111A, ssDNA, ATPase, PIP-box, telomere, DNA replication

## Abstract

Mutations in the ATRX chromatin remodeller predispose to a developmental genetic disorder and cancer, but how it safeguards genome and telomere stability remains unresolved. Here, we uncover critical dependencies for the CTC1-STN1-TEN1 (CST) complex and RAD9A-HUS1-RAD1 (9-1-1) clamp in *ATRX* deficient cells. *ATRX:CST* synthetic lethality manifests following accumulation of telomeric G-rich ssDNA, which results in telomere loss and cell death. Conversely, we attribute *ATRX:9-1-1* synthetic lethality to genome-wide ssDNA lesions, which compromise DNA replication. We further show ATRX suppresses DNA damage during replication stress by counteracting the activity of the FAM111A protease. We demonstrate that roles of ATRX in telomere maintenance and replication are genetically separable requiring its ATPase activity and PIP-box, respectively, and independently of its DAXX interaction. Collectively, functions of ATRX in suppressing toxic ssDNA lesions are context-dependent and are key to global DNA replication and telomere integrity.

---

Mutations in DNA repair genes are associated with developmental syndromes and cancer predisposition, implying that endogenous DNA damage can drive human disease. One source of endogenous damage is replication stress, which can arise within repetitive DNA that has the potential to form DNA secondary structures, transcription–replication conflicts, abasic sites, ribonucleotide misincorporation or DNA–protein crosslinks that present significant obstacles for DNA replication^1^. Replication stress can manifest in distinct regions of the genome and poses a threat to genome and epigenome maintenance, and thus, cell proliferation and identity.

Cells have evolved responses to replication stress critical for maintaining genomic integrity, which encompass the checkpoint and DNA damage tolerance pathways. DNA replication fork stalling results in stretches of ssDNA, which are coated by RPA. RPA-coated ssDNA serves as a platform to recruit and activate the ATR-ATRIP kinase complex^2,3^. ATR activation at stalled forks relies on two distinct pathways, the first involving TOPBP1 and the second involving ETAA1^4–6^. TOPBP1-mediated ATR activation depends on the heterotrimeric clamp complex RAD9A-HUS1-RAD1 (9-1-1), loaded by the alternative clamp loader RAD17-RFC onto 5’-ss/dsDNA junctions^4^. ETAA1-mediated ATR activation involves binding RPA-ssDNA directly^5,6^. ATR kinase signalling, resulting in CHK1 phosphorylation^7,8^, orchestrates the response to replication stress by enforcing the S/G2 checkpoint, suppressing origin firing and promoting fork stability and restart^9^.

Telomeres are intrinsically difficult to replicate and are a source of endogenous replication stress due to their G-rich nature and propensity to form secondary structures^10^. To aid with resolution of telomeric structures, different factors cooperate to prevent fork stalling. The CTC1-STN1-TEN1 (CST) complex binds G-rich telomeric ssDNA^11^, ensures timely disengagement of telomerase from telomeres^12,13^ and recruits DNA polymerase-α/primase to perform C-strand fill-in synthesis^14–16^. CST also plays a role in genome-wide DNA replication by stabilising stalled forks under stress and promoting their restart^17–22^ as well as in double-strand break (DSB) repair by counteracting end resection^23^.

Germline mutations in the *ATRX* gene are associated with a rare genetic disorder known as ATR-X syndrome, characterised by intellectual disability and developmental abnormalities^24^. ATRX somatic mutations are also particularly prevalent in specific types of tumours (detected in ∼20% of sarcomas, ∼40% of certain brain tumours, ∼15% pancreatic neuroendocrine tumours)^25,26^, making it one of the most mutated genes in cancer. ATRX, a chromatin remodeller of the SWI/SNF family, is known to form a complex with the histone H3.3 chaperone DAXX^27–29^, which facilitates histone H3.3 deposition at a wide range of repetitive heterochromatic genomic regions, many of which are enriched in putative G-quadruplex (G4)-DNA forming sequences, including telomeres^30–34^. ATRX-DAXX also binds and deposits H3.3 at euchromatic regulatory elements such as CpG islands^35^ and suppresses spurious transcription and replication stress within heterochromatin^36–39^. To date, the reported roles of ATRX-DAXX complex as an H3.3 chaperone, during metabolism of G4-DNA^34^ and during homologous recombination^40^ require ATRX’s chromatin binding ADD/PHD domain, interaction with DAXX, interaction with PCNA (PIP-box) and its ATPase activity. However, the underlying molecular mechanisms by which ATRX safeguards genome and telomere stability remain to be elucidated.

Here, we uncover unappreciated synthetic lethal genetic interactions between ATRX deficiency and two key ssDNA metabolism complexes, CST and 9-1-1, highlighting critical roles for ATRX in averting toxic ssDNA accumulation across the genome. Whereas the loss of *ATRX:CST* creates severe telomere instability and telomere loss following telomeric G-rich ssDNA accrual, the loss of *ATRX:9-1-1* leads to replication fork stalling and collapse that compromise genome-wide replication. We show that ATRX limits replication stress by preventing the engagement of the FAM111A protease with active replication forks. Unexpectedly, we demonstrate that the ability of ATRX to suppress accumulation of toxic ssDNA lesions at telomeres and genome-wide is independent of DAXX and HP1a binding, but requires its ATPase function and PCNA interaction motif, respectively. Our findings reveal genetically separable functions for ATRX in preventing the accumulation of toxic ssDNA lesions, which are essential for global DNA replication, telomere integrity and cell survival.

## Results

### ssDNA metabolism complexes protect cells from ATRX loss

To interrogate the contribution of ATRX to genome stability maintenance, we sought to identify novel genetic interactions with *ATRX* through genome-scale CRISPR/Cas9 screens. To this end, we developed isogenic constitutive *ATRX* knockout (KO) in telomerase-positive chronic myelogenous leukaemia-derived diploid eHAP cells harbouring an integrated doxycycline-inducible Cas9 (iCas9), which retained proliferation capacity comparable to that of the parental (WT) cells (**Fig. 1a,b and Extended Data Fig. 1a,b**). To define the genetic interaction network surrounding *ATRX* in human cells, we transduced the isogenic parental and *ATRX*-null eHAP cells with the lentiviral Brunello single-guide (sg) RNA library (**Fig. 1c**). We analysed dropout of sgRNA counts with the MAGeCK algorithm, comparing *ATRX*-null relative to the parental cells at two timepoints, day 6 (early) and day 16 (late) after addition of doxycycline (**Fig. 1c-e**). Gene Ontology (GO) analyses of dropout genes highlighted enrichment of several complexes involved in DNA replication and telomere metabolism (**Fig. 1f**). Genes comprising the replication checkpoint clamp 9-1-1 and unique clamp loader RAD17-RFC subunits scored as top dropout hits at the early timepoint (**Fig. 1d,f**). Additional members of the replication checkpoint pathway such as *ATR*-*ATRIP*, the 9-1-1 interactor *RHINO*^41^ and the alternative ATR activator *ETAA1*^6^ also emerge as statistically significant hits albeit with lower ranking (**Fig. 1d,e, Supplementary Table 1**). In contrast, all the CST complex subunits *CTC1, STN1* and *TEN1* scored as top hits, but only at the late timepoint (**Fig. 1e,f**). We chose to explore in detail the 9-1-1 checkpoint clamp and the CST complex, the most significant complexes at both timepoints (**Fig. 1f**), given their robust dropout and their links to the recognition and/or metabolism of ssDNA.

**Fig. 1:**
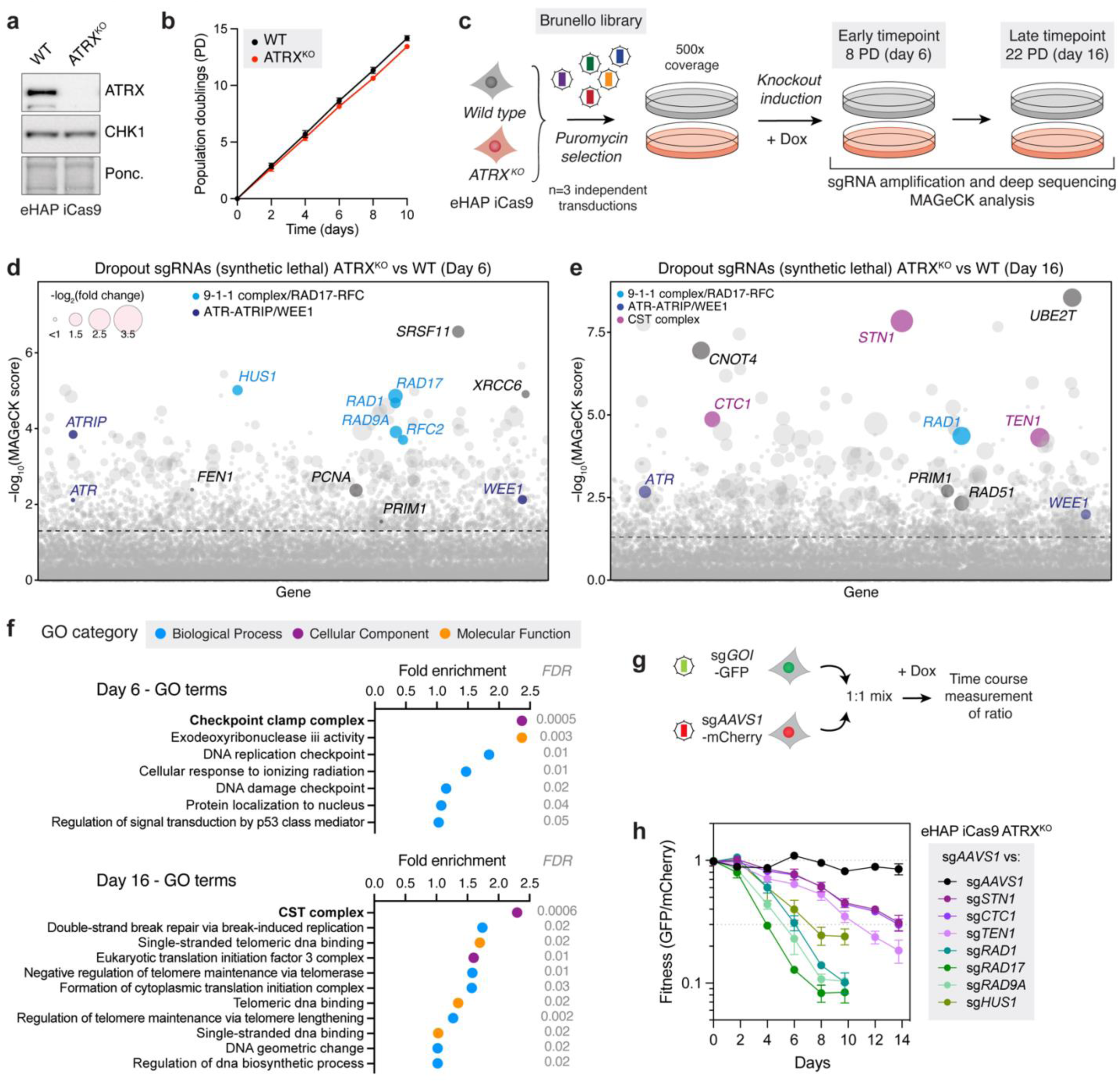
ssDNA metabolism complexes protect cells from ATRX loss. **(a)** Immunoblot of whole cell extracts (WCE) from eHAP iCas9 WT or *ATRX*-null cells, probed for ATRX. Total CHK1 and Ponceau stain used as loading control. **(b)** Growth curve in WT and *ATRX*-null eHAP iCas9 in unchallenged conditions. Data are mean ± SD (n=3 biological replicates). **(c)** Schematic of screen pipeline. **(d-e)** Bubble plot illustrating the dropout hits resulting from MAGeCK analysis at day 6 (d) and day 16 (e). Bubble size is proportional to the log2(fold change). Genes are distributed on the x axis in alphabetical order. For full hit lists, refer to Supplementary Table 1. **(f)** GO pathway enrichment of top 50 synthetic lethal genes (day 6) or top 100 synthetic lethal genes (day 16). **(g)** Schematic of two-colour competitive growth assays. **(h)** Competitive growth assays in eHAP iCas9 *ATRX*-null cells transduced with virus expressing the indicated sgRNAs against genes of interest (GOI) as in (g). Data are mean ± SEM (n=3 biological replicates).

To further validate the genetic interaction between *ATRX* and the 9-1-1 DNA replication checkpoint clamp observed at the early timepoint, we carried out live-cell imaging 96 hours after Cas9 induction (**Extended Data Fig. 1c**). Inducible knockout of the 9-1-1 clamp subunits *RAD9A*, *HUS1*, *RAD1* and the alternative clamp loader subunit *RAD17* showed a synergistic growth defect in eHAP *ATRX*-null cells, whereas the wild-type tolerated 9-1-1/RAD17 loss and achieved confluency (**Extended Data Fig. 1d-i**). To confirm the genetic dependency between *ATRX* and the CST complex observed at the late timepoint, we carried out clonogenic survival assays in which we inducibly ablated the CST complex subunits *STN1* and *CTC1* in parental or *ATRX*-null eHAP iCas9 cell lines. Loss of *CTC1* and *STN1* showed a very modest non-significant clonogenic survival reduction in combination with *ATRX* loss at day 10 after Cas9 induction (**Extended Data Fig. 1j-l**). However, a significant synergistic growth defect in clonogenic survival was observed when CST subunits were inducibly knocked out in *ATRX*-null cells for 14 days (**Extended Data Fig. 1l-n**). We next developed isogenic constitutive ATRX KO pairs in a second cell line harbouring an integrated iCas9, the non-small-cell lung cancer line NCI-H460, where inducible knockout of *RAD1* and *STN1* corroborated the pronounced synergistic cell killing with ATRX loss in clonogenic survival (**Extended Data Fig. 2a-e**).

To determine whether the time dependency of the two ssDNA metabolism complexes could reflect a dichotomy in the underlying biology, we undertook competitive growth assays with time-resolved imaging in eHAP and NCI-H460 iCas9 *ATRX*-null cells with sgRNAs targeting the safe-harbour locus *AAVS1* and *CTC1*, *STN1*, *TEN1*, *RAD9A*, *HUS1*, *RAD1* and *RAD17* (**Fig. 1g**). In unchallenged conditions and upon Cas9 induction, sgRNAs targeting 9-1-1/RAD17 caused a significant growth disadvantage in eHAP *ATRX*-null cells (ratio<0.3 over those targeting *AAVS1*) between days 4-7, whereas sgRNAs targeting CST only reached that level of growth inhibition between days 11-14 (**Fig. 1h**). This time difference was less marked in NCI-H460 *ATRX*-null cells, but both complexes still exhibited distinct dropout timing, with sgRNAs targeting 9-1-1/RAD17 depleting between days 6-9 and sgRNAs targeting CST on days 10-12 (**Extended Data Fig. 2f**). This raised the possibility that 9-1-1/RAD17 and CST complex activities protect against distinct aspects of ATRX deficiency.

### G-rich telomeric ssDNA is responsible for ATRX:CST genetic interaction

We next set out to understand whether the synergistic genetic interaction between *ATRX* and CST was driven by a defined genomic context that accumulates ssDNA over time upon CST loss. While CST functions in genome-wide DNA replication beyond telomere maintenance^17–21^, potential fork stalling events elsewhere are mitigated by dormant origin firing, whereas defects in telomere replication are likely to progressively accumulate over population doublings due to infrequent origin activation within telomeres^10,42^. Given the significant dropout of all CST complex subunits in *ATRX*-null cells only emerged at the late timepoint of the screen (**Fig. 1e,f,h**), we hypothesised that stable knockouts clones of CST would have high levels of telomeric ssDNA and therefore elicit the synthetic lethal phenotype earlier. To test this, we generated constitutive CST knockout cell lines, which are tolerated in eHAP cells (*STN1*-null and *CTC1*-null) (**Extended Data Fig. 3a**). As predicted^14–16^, *STN1*-null and *CTC1*-null cells accumulated G-rich telomeric ssDNA (**Fig. 2a**). IncuCyte imaging of *STN1*-null and *CTC1*-null cells transiently transfected with synthetic CRISPR RNA (crRNA) targeting *ATRX* in combination with unmodified transactivating crRNA (tracrRNA) following Cas9 expression showed exquisite sensitivity and rapid loss of viability at early time points, which prevented DKOs from achieving confluency (**Fig. 2b,c and Extended Data Fig. 3b,c**). These data suggest that the accumulation of G-rich telomeric ssDNA following loss of the CST complex place a critical dependence on ATRX.

**Fig. 2:**
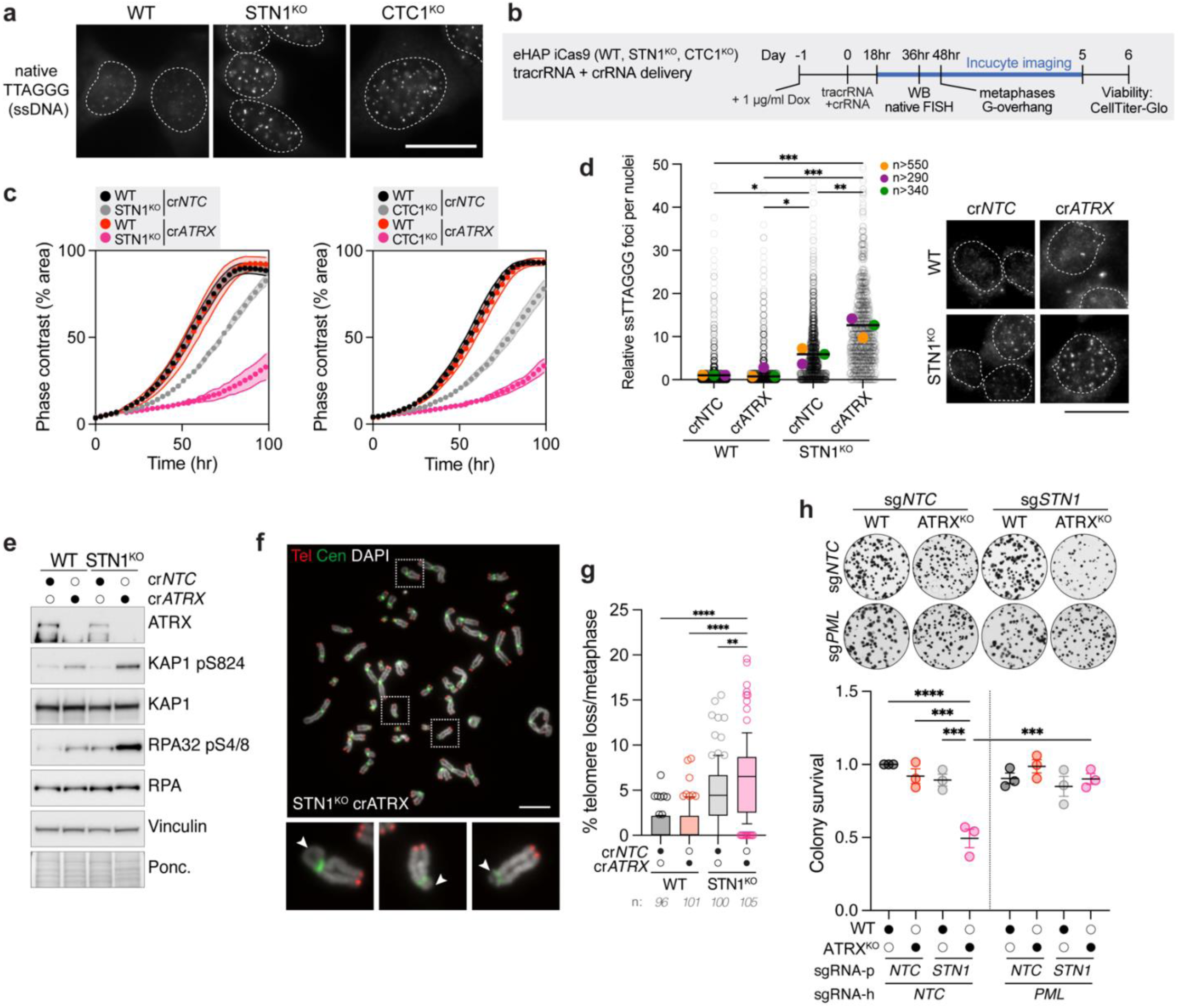
G-rich telomeric ssDNA is responsible for ATRX:CST genetic interaction. **(a)** Representative ssTTAGGG micrographs showing increased telomere G-rich ssDNA accumulation in the indicated cell lines. Scale bar: 20 µm. **(b)** Schematic of pipeline for inducible knockout transfection and downstream assays. **(c)** Proliferation curves for eHAP iCas9 WT, *STN1*-null or *CTC1*-null cells seeded 18 hours after transfecting with the indicated crRNA+tracrRNA in the presence of Dox. Data are mean ± SD of 6 technical replicates and is representative of n=4 biological replicates. **(d)** Left: quantification of ssTTAGGG foci in WT or *STN1*-null eHAP iCas9 cells 36 hours after transfecting with the indicated crRNA+tracrRNA in the presence of Dox. Cells (n, each represented as grey dot) and mean (coloured dots) of n=3 biological replicates (statistics on means, one-way ANOVA (*p<0.05, **p<0.01, ***p<0.001)). Right: representative ssTTAGGG micrographs. Scale bar: 20 µm. **(e)** Immunoblot of WCE from cells treated as in (d). Vinculin and Ponceau stain used as loading controls. **(f)** Representative images of telomere and centromere FISH in DAPI-stained metaphase spreads. Scale bar: 10 µm. **(g)** Quantification of the percent of telomeres with no signal in WT or *STN1*-null eHAP iCas9 cells 48 hours after transfecting with the indicated crRNA+tracrRNA in the presence of Dox. Number of metaphases (n) pooled from n=3 biological replicates (statistics on pooled data, one-way ANOVA (**p<0.01, ****p<0.0001)). **(h)** Top: Representative clonogenic survival assay in WT or *ATRX*-null eHAP iCas9 cells transduced with *NTC* or *STN1* sgRNA-puromycin and subsequently with *NTC* or *PML* sgRNA-hygromycin. Cells were seeded for colony formation in technical triplicate 8 days after starting Dox. Well diameter, 16 mm. Bottom: Quantification of clonogenic survival assays. Data are mean ± SEM, normalised to WT sg*NTC/NTC* cells (n=3 biological replicates, one-way ANOVA (***p<0.001, ****p<0.0001)).

Apart from the physiological G-overhang, telomeric ssDNA signal is associated with pathological circumstances such as under-replicated DNA, ssDNA gaps, or persistent replication intermediates^43,44^. We therefore systematically analysed single-stranded C-rich telomeric DNA (ssCCCTAA) and G-rich telomeric DNA (ssTTAGGG) in constitutive *STN1*-null cells. As expected, STN1^KO^ cells accumulated G-rich telomeric ssDNA foci (**Fig. 2a,d**), but did not significantly accumulate C-rich telomeric ssDNA (**Extended Data Fig. 3d**). Acute depletion of *ATRX* with a crRNA significantly enhanced the ssTTAGGG foci levels 36 hours post-transfection when compared to parental *STN1*-null cells (**Fig. 2d**). Telomeric ssDNA was not present in the form of G-or C-circles that would represent a substrate for phi29 polymerase, unlike the by-products formed during alternative lengthening of telomeres (ALT) (**Extended Data Fig. 3e**), suggesting the *ATRX:CST* genetic interaction cannot simply be explained by the previously described role of CST in promoting extrachromosomal C-circles in ALT cells^45,46^.

We next set out to test whether ATRX is required for maintenance of the G-overhang. *STN1*-null cells exhibited a ∼6.5-fold increase in G-overhang compared to parental eHAP cells (**Extended Data Fig. 3f**). After 48 hours of transfection with *ATRX* crRNA, the G-overhang did not change significantly (**Extended Data Fig. 3f**). We reasoned that in the absence of CST-mediated fill-in, if the hyper-extended G-overhang was responsible for the *ATRX:CST* genetic interaction, loss of G-strand extension by the action of telomerase would attenuate the lethality. To test this hypothesis, we abolished telomerase catalytic activity by selective chemical inhibition with BIBR1532 in NCI-H460 cells (**Extended Data Fig. 3g**). Consistent with the lack of a significant change in G-overhang length between the *STN1*-KO or cr*ATRX*-*STN1*-DKO cells, the *ATRX:STN1* synthetic genetic interaction was independent of telomerase catalytic activity (**Extended Data Fig. 3h**). These data suggest that the accumulation of ssTTAGGG that occurs after acute ATRX loss in *STN1*-null cells may represent interstitial single-stranded gaps or single-stranded internal loops that are not terminal, and independent of telomere lengthening by telomerase.

Interestingly, we noticed that *ATRX* deletion in *STN1*-null cells sparked a rapid and sustained increase in DNA damage signalling associated with DSBs, including phosphorylation of RPA32 Ser4/8, RPA32 Ser33, KAP1 Ser824 and H2AX Ser139 (**Fig. 2e and Extended Data Fig. 3i-k**). To determine how telomere integrity was affected by the loss of *ATRX* in CST deficient cells, we examined telomere FISH-stained metaphase spreads for telomere fragility, heterogeneity and loss. While acute ATRX loss did not affect telomere fragility or heterogeneity in *STN1*-null cells, it resulted in a significant increase in signal-free ends indicative of telomere loss, with an average of >5 telomeres completely devoid of detectable FISH signal per metaphase (**Fig. 2f,g and Extended Data Fig. 3l,m**). Cells lacking both STN1 and ATRX also presented with elevated levels of micronuclei when compared to the single KOs, with ∼40% of cells presenting with at least one micronucleus per primary nucleus and about a third of these with two or more micronuclei (**Extended Data Fig. 3n**).

Telomeres in *ATRX*-deficient cells were shown to exhibit increased interactions with PML bodies^47^. In normal human cells, PML nuclear bodies associate with repair factors such as BLM and MRE11 into discrete foci and few telomeres, preferentially when they are damaged^48–50^. In ALT cells, PML recruits the BLM complex^43^ responsible for the generation of single-stranded telomere intermediates^46,51^. To test whether PML could suppress the generation of telomeric G-rich ssDNA in eHAP sg*STN1-ATRX-*DKO cells and induction of the damage response, we leveraged the prolonged inducible system with sgRNAs against *STN1* and/or *PML* in parental or *ATRX*-null cells (**Extended Data Fig. 3o**). The phosphorylation of RPA32 Ser33 and H2AX Ser139 observed in sg*STN1-ATRX*-DKO cells was significantly reduced upon loss of PML, implying that PML is responsible for DNA damage accumulation in sg*STN1-ATRX*-DKO cells (**Extended Data Fig. 3p,q**). PML loss also completely abolished G-rich ssDNA present in constitutive *ATRX*-null cells, whereas only partially reduced G-rich ssDNA present in inducible sg*STN1* cells, implicating PML in ssDNA generation upon ATRX loss and a subset of ssDNA generated in CST deficient cells (**Extended Data Fig. 3r**). G-rich ssDNA in the sg*STN1-PML-ATRX* triple KO was comparable to that in the sg*STN1-PML*-DKO, suggesting the signal present in sg*STN1-PML*-DKO might represent G-overhangs, which supports the notion that those are PML-independent. We next asked whether PML could drive cell death in the context of the *ATRX:STN1* genetic interaction. Strikingly, the synergistic growth defect of sg*STN1-ATRX*-DKO cells was entirely rescued by loss of PML (**Fig. 2h**). We conclude that ATRX ensures telomere integrity through inhibition of PML-dependent telomeric ssDNA formation, and this becomes critical for cell survival when G-rich ssDNA accumulates upon prolonged absence of CST.

### 9-1-1 cooperates with ATRX to suppress global replication fork collapse

Loss of 9-1-1 complex resulted in a loss of viability of *ATRX*-null cells, as observed and validated in an early timepoint of the screen (**Fig. 1d,f,h**). To understand how the combined loss of ATRX and 9-1-1/RAD17 promotes cell death, we examined DNA damage signalling in cells expressing *NTC* or *RAD17*-targeting sgRNAs 72 hours after Cas9 activation. Loss of *RAD17* in *ATRX*-null cells resulted in robust activation of DNA damage signalling, including phosphorylation of CHK1 Ser345, RPA32 Ser4/8 and H2AX Ser139 (**Fig. 3a**). We assume that CHK1 phosphorylation is achieved in 9-1-1 deficient cells devoid of ATR signalling by crosstalk between PIKKs (ATM or DNA-PK), as reported previously^52,53^. Additionally, activation of apoptosis was evident from PARP1 and Caspase-3 cleavage (**Fig. 3a**). *ATRX*-null cells or ablation of *RAD17* in parental WT cells had only mild effects on the cell cycle, whereas ablation of *RAD17* in *ATRX*-null cells resulted in alterations in S-phase and an accumulation of a sub-G1 population, consistent with apoptosis (**Fig. 3b-d**). Notably, in sg*RAD17*-*ATRX* double KO (DKO) cells, ∼10% of S-phase cells failed to incorporate the nucleotide analogue EdU suggestive of compromised DNA replication (**Fig. 3c-e**).

**Fig. 3:**
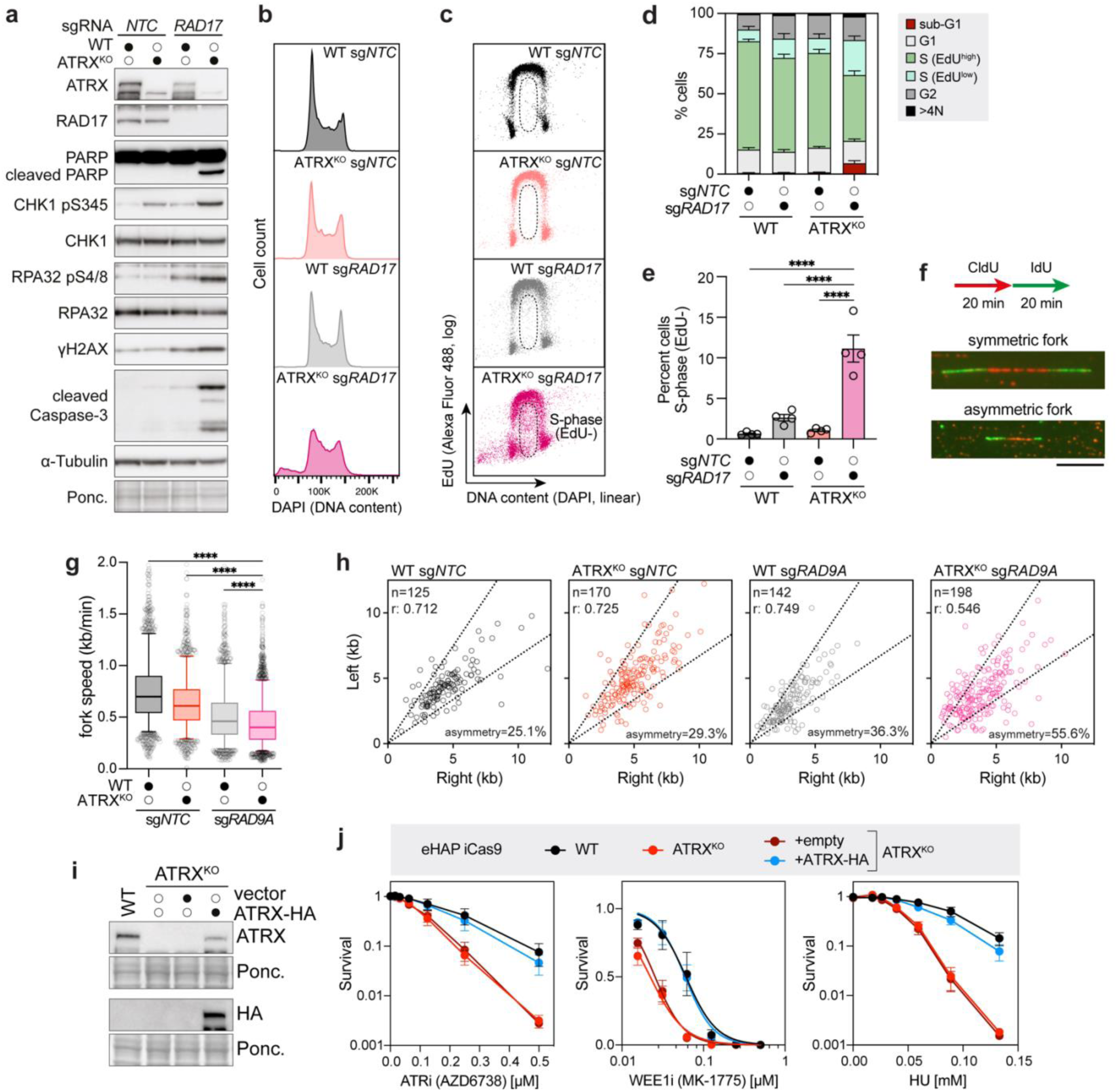
9-1-1 cooperates with ATRX to suppress global replication fork collapse. **(a)** Immunoblot of WCE from WT or *ATRX*-null eHAP iCas9 cells transduced with *NTC* or *RAD17* sgRNA following 72 h Dox. Alpha-tubulin and Ponceau stain used as loading controls. **(b)** Representative FACS DNA content analysis of eHAP iCas9 cells treated as in (a). **(c)** Representative FACS EdU/DNA content plots of eHAP iCas9 cells treated as in (a) and pulsed with EdU. **(d)** Quantification of % of cells in different cell-cycle stages. WT or *ATRX*-null eHAP iCas9 cells were transduced with *NTC* or *RAD17* sgRNA following 72 h Dox and pulsed with EdU. Data are mean ± SEM (n=4 biological replicates). **(e)** Quantification of the S-phase (EdU-) population as gated in (c). Data are mean ± SEM (n=4 biological replicates, one-way ANOVA (****p<0.0001)). **(f)** Experimental setup and representative images of DNA fibers. Scale bar: 10 µm. **(g)** Distribution of replication fork speeds of DNA fibers prepared as in (f). Data are pooled from n=3 biological replicates. **(h)** Scatterplot of DNA fibers fork asymmetry prepared as in (f). Data are pooled from n=3 biological replicates. Note that one or two data points are outside axis limits in *ATRX*-null sg*NTC* or *ATRX*-null sg*RAD9A*, respectively. n, number of bidirectional DNA fibers analysed, r, Pearson coefficient. **(i)** Immunoblot of WCE from WT or *ATRX*-null eHAP iCas9 cells stably expressing HA-tagged ATRX cDNA. Ponceau stain used as loading control. **(j)** eHAP iCas9 cells as in (i) were treated continuously for 5 days with the indicated doses of ATRi (AZD6738), WEE1i (MK-1775) or hydroxyurea (HU), and viability was determined using CellTiter-Glo. Data are mean ± SEM, normalised to untreated cells (n=3 [ATRi and WEE1i] or n=5 [HU] biological replicates).

To examine DNA replication directly, we monitored the progression of individual replisomes by measuring fork extension rates and symmetry of bidirectional forks using the DNA fibre assay. To this end, we sequentially pulse-labelled active replication forks with CldU and IdU nucleotide analogues and visualised the labelled replication forks by immunofluorescence (**Fig. 3f**). *ATRX* or *9-1-1* single knockouts presented with reduced replication fork extension rates and a modest increase in fork asymmetry, which is suggestive of fork stalling and dormant origin activation. Combined loss of ATRX and 9-1-1 resulted in a further attenuation in fork speed when compared with parental and single KOs (**Fig. 3g**). Consistent with the significant population of cells failing to incorporate EdU (**Fig. 3c-e**), >50% of newly activated replication origins exhibited fork asymmetry in sg*RAD9A*-*ATRX* DKO cells (**Fig. 3h and Extended Data Fig. 4a**). *ATRX*-null cells showed heightened sensitivity to chemical inhibition of the DNA replication checkpoint via an ATR inhibitor (ATRi) or the G2/M checkpoint via a WEE1 inhibitor^54^ (WEE1i), as well as to DNA replication perturbation following hydroxyurea (HU), in both eHAP and NCI-H460 cells (**Fig. 3i,j and Extended Data Fig. 4b,c**). Taken together, these findings suggest that ATRX is required to prevent the accumulation of endogenous replication-borne toxic lesions in S phase and the observed synthetic lethality with 9-1-1/RAD17 stems from global replication fork stalling and/or collapse, which triggers apoptosis.

### ATRX restrains the engagement of FAM111A at active replication forks

Based on the critical role of ATRX in conditions of replication stress in different genomic contexts, we hypothesized that ATRX may perform previously unappreciated functions in the recruitment and/or turnover of specific factors at replication forks. To explore this possibility, we performed isolation of proteins on nascent DNA (iPOND) coupled with stable isotope labelling with amino acids in cell culture (SILAC)-based quantitative mass spectrometry to compare protein occupancy at the fork between WT and *ATRX*-null cells^55^. We compared protein enrichments at active replication forks (EdU-pulse labelled nascent chromatin) and at post-replicative chromatin that was allowed to mature during a 60-minute chase. Additionally, we included a chase sample in the presence of HU to enrich for stalled replication forks (**Fig. 4a,b**). Our approach identified ATRX significantly enriched in WT eHAP cells in all conditions including active forks, stalled forks, and post-replicative chromatin (**Fig. 4c-e and Supplementary Table 2**), consistent with previous iPOND^56–58^ and nascent chromatin capture experiments^59^. Within the identified replication and chromatin associated factors, 36 proteins emerged as differentially enriched 1.4-fold average or greater in *ATRX*-null cells in both biological replicates (**Fig. 4c-f and Extended Data Fig. 5a-d**). Interestingly, the HU-treated samples uncovered many chromatin-associated factors directly involved in chromatin condensation, remodelling and/or repair including SUMO2, TRIM28, members of the ChAHP (CHD4, ADNP, HP1) and nucleosome remodelling and deacetylase (NuRD) repressor complexes, linker histone H1, histone H2A variants H2A.V, H2AX and macroH2A.1, including its histone chaperone and helicase HELLS (**Fig. 4d,f and Extended Data Fig. 5b,d**). These data suggested that nascent chromatin is most affected in the absence of ATRX upon severe fork stalling and replication stress.

**Fig. 4:**
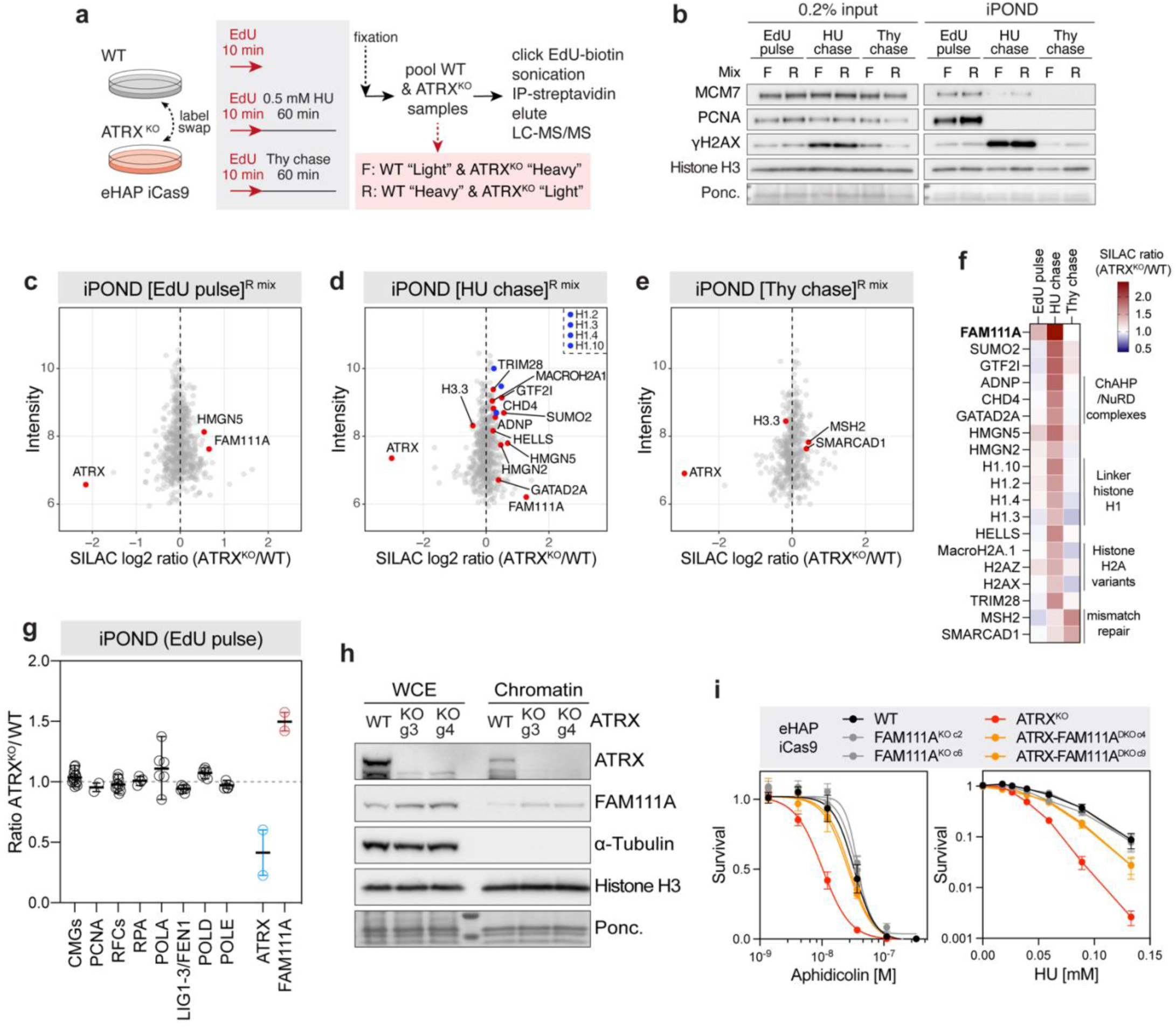
ATRX restrains the engagement of FAM111A at active replication forks. **(a)** Schematic of iPOND-SILAC-MS strategy for proteomics analysis of active and stalled replication forks, and post-replicative chromatin. The “heavy” and “light” isotope label was swapped during SILAC labelling for the two biological repeats, resulting in pooled forward (F) and reverse (R) mixes before iPOND purification. **(b)** Immunoblot of input and iPOND pull-downs. MCM7 and PCNA are controls for active forks, γH2AX is a control for damaged forks whereas Histone H3 and Ponceau stain are used as loading controls. **(c-e)** Volcano plots of the intensity versus the ATRX^KO^/WT SILAC log2 ratio of the iPOND-MS identified proteins in active replication forks (c), stalled replication forks (d) and post-replicative chromatin (e). Data points highlighted represent proteins enriched/depleted significantly changed in both biological replicates. **(f)** Heat map of ATRX^KO^/WT SILAC ratio of proteins identified as enriched in both biological replicates of iPOND-MS. **(g)** Enrichment showing the WT/ATRX^KO^ ratio of SILAC iPOND-MS identified proteins at active replication forks. For CMGs, RFCs, RPAs, replicative polymerases, and LIG1-3/FEN1 mean ± SEM was calculated from the SILAC ratio of each subunit. For ATRX and FAM111A, the SILAC ratio is used to calculate mean ± SD (n=2 biological replicates). **(h)** Immunoblot of WCE versus chromatin from WT or *ATRX*-null asynchronous eHAP cells. Alpha-tubulin was used as a soluble fraction loading control. Histone H3 was used as a chromatin fraction loading control. Data are representative of 4 independent experiments. **(i)** eHAP cells were treated continuously for five days with aphidicolin or HU at the indicated doses, and viability was determined using CellTiter-Glo. Data are mean ± SEM, normalised to untreated (n=6 (aphidicolin) or n=7 (HU) biological replicates).

The enrichment of core replisome machinery such as CMGs, PCNA, RFCs, RPA, DNA polymerases (POLD, POLA and POLE) and lagging strand processing (LIG1, LIG3 and FEN1) did not change remarkably upon ATRX loss at active replication forks (**Fig. 4g**). In contrast, FAM111A was one of the few proteins significantly enriched in *ATRX*-null cells at active replication forks in unchallenged conditions and was also detected as enriched in HU-treated conditions (**Fig. 4c,d,f,g**). FAM111A is a PCNA-interacting protease mutated *de novo* in Kenny-Caffey syndrome type 2 and Gracile Bone Dysplasia, which is known to interact with active replication forks and exhibits specific ssDNA-binding activity *in vitro*^57,59–61^. While its protease activity has been shown to protect against topoisomerase-induced DNA-protein crosslinks (DPCs)^61^, its unrestrained function, best illustrated by ectopic overexpression of wild-type or the constitutively active R569H mutant, results in toxic ssDNA formation, replicative stress and cell death^62,63^. To validate that FAM111A differentially associates with newly synthesized DNA in *ATRX*-null cells, we took advantage of the rapid cycling time of eHAP cells, with asynchronous cultures mostly in S-phase (**Fig. 3d**), and compared levels in the chromatin fraction. Consistent with iPOND-SILAC proteomics, two independent *ATRX*-null clones exhibited significant enrichment of FAM111A on chromatin (**Fig. 4h**).

To determine whether FAM111A functions during the replication stress response, we evaluated survival of cells genetically inactivated for *FAM111A* in parental and *ATRX*-null backgrounds (**Extended Data Fig. 5e**) in response to replication inhibiting drugs. Loss of FAM111A did not affect the survival after aphidicolin or HU treatment compared to wild-type cells (**Fig. 4i**). However, combined loss of *FAM111A* and *ATRX* alleviated ATRX-dependent aphidicolin and HU hypersensitisation (**Fig. 4i**). The HU resistance observed in *ATRX*-null eHAP cells upon *FAM111A* loss is consistent with previous studies in U2OS cells (ALT-positive *ATRX*-deficient)^62,63^. Importantly, *FAM111A* knockout in *ATRX*-deficient cells was also sufficient to attenuate HU-induced phosphorylation of RPA32 Ser4/8 and H2AX Ser139 (**Extended Data Fig. 5f**).

Next, we asked whether *FAM111A*-deficiency impacts the cellular response of *ATRX*-deficient cells to replication stress caused by checkpoint inhibition. Exposure to ATR or WEE1 inhibitors revealed moderately enhanced sensitivity of the *FAM111A*-null cells compared to WT control (**Extended Data Fig. 5g**). In contrast to HU or aphidicolin sensitivity, deletion of FAM111A did not suppresses the hypersensitivity of *ATRX*-null cells to ATR or WEE1 inhibitors. Consistently, deleting FAM111A failed to rescue the synthetic lethality of untreated *ATRX:9-1-1* DKO cells (**Extended Data Fig. 5h,i**). The differential impact of FAM111A loss on the survival of *ATRX*-null cells upon replication stalling or checkpoint inhibition, suggests that other factors, such as unrestrained origin firing and replication fork stabilisation/repair, contribute to the loss of viability seen in checkpoint inhibited or *ATRX:9-1-1* cells, which is FAM111A independent.

### ATRX ATPase and PIP-box suppress telomeric ssDNA and genome-wide replicative stress

ATRX is implicated in chromatin maintenance and DNA repair processes, which rely on distinct protein-protein interactions (**Fig. 5a**). The amino-terminal ATRX-DNMT3-DNMT3L (ADD) domain and the conserved LXVXL HP1-interacting motif are responsible for binding to heterochromatin (H3K9me3/H3K4me0 and HP1α, respectively)^64,65^. ATRX binding to the histone chaperone DAXX requires a minimal region of ∼30 amino acids and can be disrupted by mutation of Leu 1276^66^ (**Extended Data Fig. 6a**). A PCNA-interacting peptide (PIP)-box in ATRX located at residues 1843-1850 is required for irradiation-induced extended DNA repair synthesis^40^. Finally, the SNF2-type ATPase/helicase enzymatic activity in the carboxy-terminal domain has been proposed to facilitate histone exchanges at repetitive sequences and can be abolished by mutation of Lys 1600^67,68^.

**Fig. 5:**
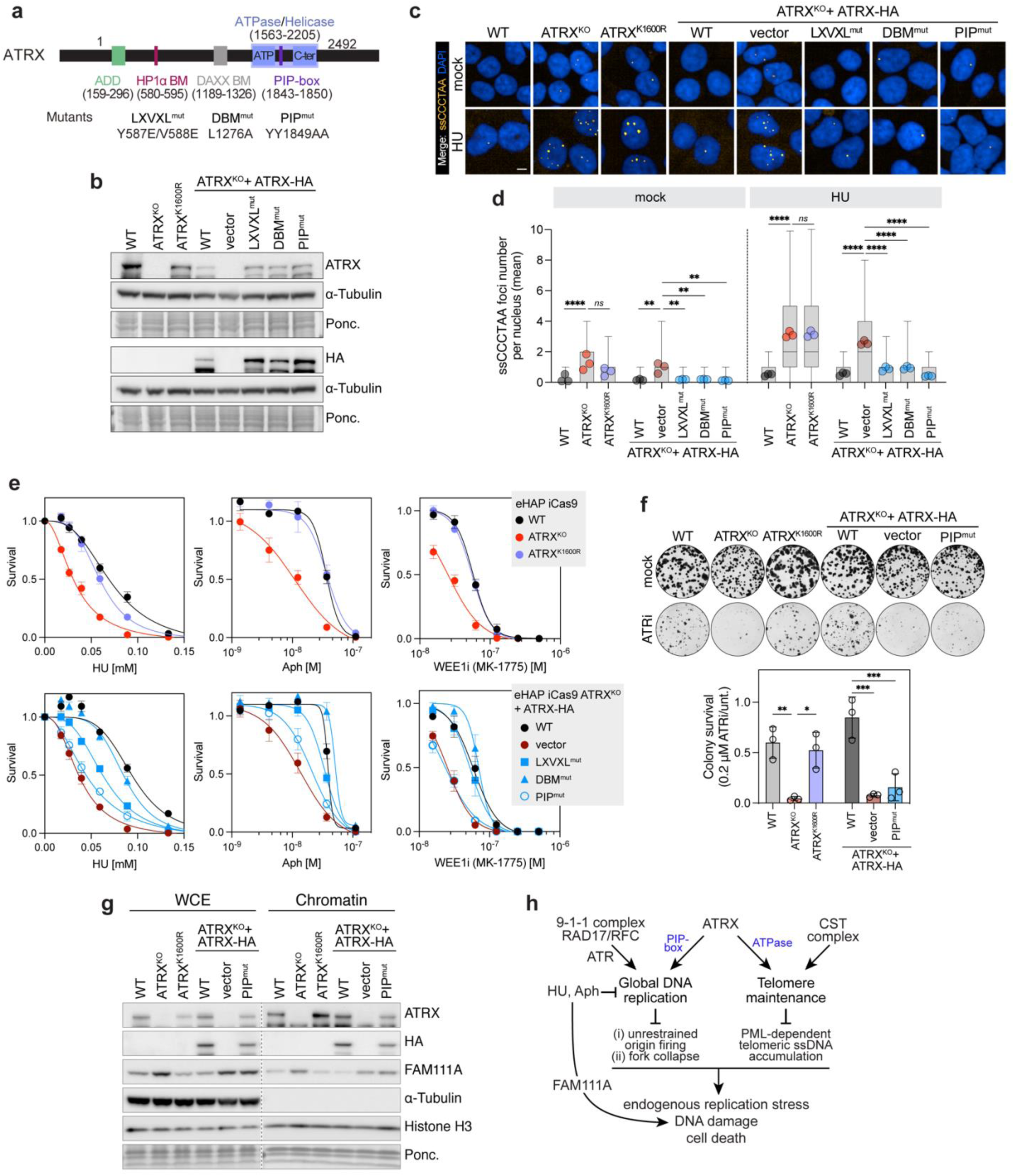
ATRX ATPase and PIP-box suppress telomeric ssDNA and genome-wide replicative stress. **(a)** Schematic of full-length ATRX and mutants used to reconstitute *ATRX*-null cells. **(b)** Immunoblot of WCE from cell lines including K1600R knockin and *ATRX*-null cells stably expressing HA-tagged *ATRX* cDNA mutants. Alpha-tubulin and Ponceau stain used as loading controls. **(c)** Representative ssCCCTAA micrographs showing increased telomere C-rich ssDNA accumulation in the indicated cell lines in the presence or absence of 0.15 mM HU. Scale bar: 10 µm. **(d)** Quantification of ssCCCTAA foci per nucleus in cells as in (c). Box plots represent individual nuclei with IQR, horizontal line indicates median, and whiskers extend to 5th and 95th percentile. Data points outside this range are not shown. Coloured dots are means of n=3 biological replicates, with a minimum average of 600 nuclei per replicate (one-way ANOVA (**p<0.01, ****p<0.0001)). **(e)** The indicated cell lines were treated continuously for 5 days with the indicated doses of HU, aphidicolin or WEE1 inhibitor MK-1775, and viability was determined using CellTiter-Glo. Data are mean ± SEM, normalised to untreated cells (n=3 [HU and WEE1i], n=4 [aphidicolin] biological replicates). **(f)** Top: Representative clonogenic survival micrographs of eHAP cells ± continuous treatment with ATR inhibitor for 5 days. Well diameter, 16 mm. Bottom: Quantification of clonogenic survival assays as indicated. Data are mean ± SD, normalised to untreated cells (n=3 biological replicates, one-way ANOVA (*p<0.05, **p<0.01, ***p<0.001). **(g)** Immunoblot of WCE versus chromatin from the indicated eHAP cells. Alpha-tubulin was used as a soluble fraction loading control. Histone H3 was used as a chromatin fraction loading control. Data are representative of 2 independent experiments. **(h)** Schematic model that illustrates how ATRX cooperates with 9-1-1 (through its PIP-box) and CST (through its ATPase) complexes to suppress toxic ssDNA and maintain global DNA replication and telomere integrity, respectively.

To determine if each of these domains in ATRX are required for protection against telomeric and global ssDNA lesions, we generated the indicated panel of point mutations (**Fig. 5a**). Complementation of *ATRX*-null cells using a C-terminal HA-tag stable expression system was achieved for wild-type and most mutants (**Fig. 3i and Fig. 5b**). However, the ATPase inactive ATRX K1600R mutation displayed severely reduced expression, which precluded its phenotypic analysis. We therefore generated a homozygous knockin cell line by directly mutating Lys 1600 by Cas9-mediated homologous recombination at the endogenous *ATRX* locus in wild-type cells (**Fig. 5b and Extended Data Fig. 6b**).

The ATRX ATPase inactive K1600R mutation significantly impaired its ability to supress the generation of telomeric ssDNA, as assessed by native FISH using both G-rich and C-rich probes (**Fig. 5c,d and Extended Data Fig. 6c**). This was most apparent after treating cells with a low dose of replication inhibitor HU but was also observed for C-rich ssDNA in unchallenged cells (**Fig. 5c,d and Extended Data Fig. 6c**). Surprisingly, ATPase inactive ATRX K1600R mutant cells were not hypersensitive to treatments that inhibit genome-wide replication such as HU or aphidicolin (**Fig. 5e and Extended Data Fig. 6d,e**). In contrast, genome-wide replication stress sensitivity of *ATRX*-null cells was recapitulated with a PIP-box mutant, which was dispensable for the generation of telomeric ssDNA (**Fig. 5c-e and Extended Data Fig. 6c-e**). Unexpectedly, the ATRX-DAXX interaction mediated by ATRX Leu 1276 residue and essential for assembly of the ATRX-DAXX H3.3 chaperone complex, was largely dispensable for supressing telomeric ssDNA or promoting cell fitness upon replication inhibition (**Fig. 5c-e and Extended Data Fig. 6c-e**). Similarly, the ATRX-HP1α interaction, mediated by the LXVXL motif, was not required to supress telomeric ssDNA and only mildly required to promote survival upon replication stress (**Fig. 5c-e and Extended Data Fig. 6c-e**).

By subjecting the mutant cell lines to clonogenic survival assays with chronic ATR inhibitor treatment, we observed a significant synergistic growth defect with ATR inhibition upon ATRX loss or with the PIP-mutant (**Fig. 5f**). Similar results were obtained using a WEE1 inhibitor, where the hypersensitivity of *ATRX*-null cells was recapitulated with the PIP-mutant but not with the ATPase inactive mutant (**Fig. 5e**). Transient transfection with synthetic crRNA targeting *RAD1* following Cas9 expression consistently showed that the ATRX PIP-box is required to protect cells against failure to activate the replication checkpoint (**Extended Data Fig. 6f**). Conversely, ATRX ATPase mutant cells displayed a growth defect in combination with *STN1* depletion (**Extended Data Fig. 6g**). In line with the notion that the PIP-mutant impacts genome-wide replication stress, chromatin engagement of FAM111A was dependent on ATRX PIP-box but independent of ATRX ATPase activity (**Fig. 5g**). Collectively, our data reveal genetically separable functions for ATRX during DNA replication and telomere maintenance independent of its roles in the ATRX-DAXX complex: the ATPase activity of ATRX is essential for restricting telomeric ssDNA formation but is dispensable for genome-wide replication, whereas the PCNA-interacting motif is essential to suppress genome-wide replication stress but is dispensable for telomere maintenance.

## Discussion

While *ATRX* is one of the most mutated genes in genetic disease and cancer, how the ATRX chromatin remodeller safeguards genome and telomere stability remains unresolved. Here, we uncover context-dependent functions of ATRX in supressing toxic ssDNA lesions and the compensatory pathways that ATRX deficient cells critically rely on to prevent genome instability and cell death (**Fig. 5h**). Our data reveal previously unappreciated intrinsic functions for ATRX in genome maintenance independent of DAXX binding and its canonical function as an H3.3 chaperone. While DAXX has been shown to suppress R-loops to maintain centromere stability and transcriptional silencing of endogenous retroviruses independent of ATRX^66,69,70^, we show here that ATRX supresses toxic ssDNA accumulation independent of DAXX. Contrary to current dogma, we show that the function of ATRX at telomeres requires its ATPase activity, but not its PIP-box motif, whereas its role during DNA replication requires its PIP-box motif but not its ATPase activity.

We identify a synthetic lethal interaction between *ATRX:*CST, linking ATRX to the suppression of PML-dependent telomeric ssDNA formation (**Fig. 2**). Normally, the CST complex limits the accumulation of G-rich ssDNA and its loss results in progressive telomeric G-rich ssDNA accrual, with the magnitude of this phenotype correlating with the severity of the *ATRX:CST* synthetic lethality. It is possible that the lengthened G-overhang in CST deficient cells assembles an extended D-loop within the telomeric loop (t-loop), a structure where the 3′ ssDNA overhang loops back and invades the double stranded telomeric repeat^71^. Based on our data, these D-loops could be ∼6.5-fold longer than in wild-type cells. We considered a role for ATRX in maintaining an extended D-loop could be analogous to previous work implicating ATRX in repair of DSBs by homologous recombination (HR), downstream of RAD51 filament assembly and the action of RAD54^40^. However, the telomeric ATRX function we uncover depends exclusively on its ATPase activity, whereas ATRX’s role during HR^40^ requires both its ATPase and PIP box activities, suggesting a distinct mechanism.

Instead, the most parsimonious explanation is that cells lacking CST accumulate internal G-rich telomeric ssDNA (C-strand gaps), corresponding to the fraction of G-rich telomeric ssDNA resistant to exonuclease treatment^72–74^, which creates the exquisite dependency on ATRX ATPase-dependent nucleosome remodelling promoting complete replication of internal telomeric DNA tracts. This model is consistent with several observations: ATRX protects CST deficient cells from telomere catastrophe independent of telomerase and PML loss completely rescues the *ATRX:STN1* lethality while not affecting the majority of STN1-dependent G-rich telomeric ssDNA, which presumably corresponds to the G-overhang. We speculate that interstitial telomeric ssDNA may invade neighbouring DNA creating loops that are subject to telomere trimming within PML nuclear bodies. Alternatively, fork stalling when encountering a gapped substrate may localise to PML bodies and result in one-ended DSBs resulting in telomere loss. Previous reports demonstrated that CST complex activity, via POLA-PRIM, initiates the synthesis of complementary C-rich lagging strands in exposed gap or G4-DNA structures^74^. Our data suggests telomeric ssDNA generation is solely dependent on ATRX ATPase function and decoupled from the H3.3 chaperone complex, unlike suppression of G4-DNA induced stress where both ATPase and DAXX-interaction are essential^34^. Together, these findings support a model in which CST and ATRX act in parallel to maintain telomere integrity, with ATRX averting catastrophic processing within PML bodies when CST is absent.

We define a synthetic lethal interaction between *ATRX:9-1-1*, which is recapitulated with an ATR inhibitor, implicating the ATRX PIP-box in protecting against endogenous lesions that critically rely on the replication checkpoint clamp 9-1-1. Indeed, our data suggest that loss of viability of *ATRX:9-1-1* DKO cells results from spontaneous replication fork collapse, which culminates in apoptosis (**Fig. 3**). Although the precise molecular nature of genome-wide DNA stalled structures remains to be determined, our data lead us to suggest that ATRX PIP-box function escorting the replisome is key to prevent 5’-ss/dsDNA junctions that recruit 9-1-1. Such a role would be consistent with observations showing that 9-1-1 complex protects from replication fork stalling by shielding Okazaki fragments downstream of the replicative block from endonucleolytic attack^75^. While the endogenous stalling lesion in *ATRX*-null cells could be DNA secondary structures, both ATRX ATPase activity and DAXX-interaction are required to safeguard from G4-DNA induced stress^34^. Our data suggests that both these activities are dispensable for protection against replication perturbation and S-phase checkpoint inhibition, while ATRX PIP-box is required, and therefore endogenous genome-wide lesions are likely independent from G4-DNA.

Our data shows ATRX prevents toxic lesions during DNA replication perturbation independently of ATRX ATPase activity, ATRX-DAXX and ATRX-HP1α interactions, mediated by ATRX L1276 residue and LXVXL motif, respectively (**Fig. 5**). This establishes the existence of a non-canonical function of ATRX that would escort the replisome, independent of H3.3 deposition, but dependent on its interaction with PCNA. Indeed, our experiments establish that ATRX counteracts chromatin engagement of FAM111A, particularly at active or stalled replication forks and independently of ATRX ATPase activity (**Fig. 4 and 5**). We further show that the ssDNA inducer FAM111A is responsible for damage accumulation at stalled replication forks in *ATRX*-deficient cells and is a key determinant of sensitivity to replication inhibition. Notably, FAM111A expression appears elevated in ATRX deficiency (**Fig. 4h and 5g**), which may reflect a potential link between its chromatin association and protein stability as well as indirect transcriptional regulation^76^. FAM111A overexpression has been reported to undergo auto-cleavage, triggering its protease activity, which may increase ssDNA at forks via cleavage of replisome components such as RFC1^62^. However, this was not observed in the presence of endogenous levels of FAM111A protease. To date, efforts to identify targets of FAM111A beyond its role as a viral restriction factor have revealed FAM111A itself and nuclear pore-associated factors as candidate substrates in unchallenged cells, and TOP1 and PARP1 in cells treated with topoisomerase and PARP inhibitors^61–63,77^. The increased FAM111A loading at replication forks we observe in *ATRX*-null cells could still be in an auto-inhibited state from its own N-terminal region^78^ and its release could be primed by exogenous damage. While ATRX function could hinder FAM111A fork accessibility by limiting the presence of ssDNA, FAM111A could also act on endogenous ssDNA lesions subsequently creating vulnerabilities in *ATRX* deficient cells that require unique ssDNA metabolism complexes for their repair.

## Acknowledgments

We thank members of the Boulton lab for their input and the Francis Crick Institute’s Genomics, Advanced Light Microscopy, Making Lab, Flow Cytometry, Bioinformatics & Biostatistics, Cell Science and High Throughput Screening science technology platforms. We also thank Travis Stracker for critical reading of the manuscript. S.S-B. was supported by an EMBO Long Term Fellowship (ALTF 707-2019) and by the European Union’s Horizon 2020 research and innovation programme under the Marie Sklodowska-Curie grant agreement No 886577. G.H. is supported by the Radiation Research Unit at the Cancer Research UK City of London Centre Award (C7893/A28990). Work in the S.J.B. lab is supported by the Francis Crick Institute (CC2057), European Research Council Advanced Investigator grants (TelMetab, ChrEndProt), a Wellcome Trust Senior Investigator Award, and CRUK RadNet City of London.

## Author contributions

Conceptualization: S.S-B. and S.J.B. Methodology: S.S-B., M.M., Z.M., T.H.S., S.H., P.K. Investigation: S.S-B., M.M., T.T., Z.M., S.L., T.H.S., A.I.I., G.H., P.R., R.M., P.K. Formal analysis: S.S-B., H.P., S.H. Resources: M.H. Data curation: S.S-B., S.H. Visualization: S.S-B. Project administration: S.S-B. and S.J.B. Funding acquisition: S.J.B. Supervision: S.J.B. Writing-original draft: S.S-B. Writing – review & editing: S.S-B., S.J.B.

## Methods

### Bacterial strains

Stbl3 *E. coli* strain (genotype: F–mcrB mrr hsdS20(rB–, mB–) recA13 supE44 ara-14 galK2 lacY1 proA2 rpsL20(StrR) xyl-5 λ–leu mtl-1) was transformed with piggybac or lentiviral mammalian expression plasmids and grown in Luria Broth at 30°C in the presence of ampicillin (100 ug/mL).

### Cell lines

The human chronic myelogenous leukemia-derived cell line eHAP (haploid purchased from Horizon Discovery albeit stable diploid cells used throughout) and the human lung cancer cell line NCI-H460 (purchased from ATCC) were further modified by a lentiviral integration of doxycycline-inducible Cas9 nuclease (iCas9) with an Edit-R inducible lentiviral Cas9 vector (Horizon Discovery)^79^. eHAP iCas9 and NCI-H460 iCas9 (parental WT cell lines) were derived as single clones from the pool of transduced cells after blasticidin selection (8 µg/mL for eHAP, 2.5 µg/mL for NCI-H460) and were chosen based on a high cutting efficiency and negligible leakiness using a BFP/GFP reporter assay (Addgene #67980). eHAP iCas9 parental cell line (and where indicated, *ATRX* KO line) was used for the generation of *ATRX*, *STN1*, *CTC1* and *FAM111A* full knockouts. eHAP cell lines were grown in Iscove′s modified Dulbecco′s medium (IMDM) with 10% Tet-free FBS and 1% penicillin/streptomycin. NCI-H460 iCas9 cells were used for the generation of *ATRX* full knockouts. NCI-H460 cell lines were grown in Roswell Park Memorial Institute (RPMI)-1640 medium with 10% Tet-free FBS and 1% penicillin/streptomycin. All cell lines were grown at 37°C in 5% CO_2_.

### Plasmids

ATRX-HA was introduced into pDONR221 (Invitrogen) by amplification from IF-GFP-ATRX (Addgene #45444) using primers attb1-ATRX-F: GGGGACAAGTTTGTACAAAAAAGCAGGCTTCATGACCGCTGAGCCCATG and attb2-ATRX-R: GGGGACCACTTTGTACAAGAAAGCTGGGTATCAAGCGTAATCTGGAACATCG.

Mutant variants were introduced by Q5 site directed mutagenesis (NEB) in pDONR221, according to manufacturer’s instructions (oligonucleotides are listed in Supplementary Table 4). Mammalian expression vectors were made in a piggyBac (PB) transposon delivery system PB-EF1α-DEST-IRES-Puro. ATRX-HA was cloned into PB-EF1α-DEST-IRES-Puro using the Gateway technology (Thermo Fisher Scientific) according to the manufacturer’s protocol. Expression constructs were introduced into eHAP iCas9 ATRX-KO cells by co-transfection with Super PiggyBac Transposase (PB200A-1, System Biosciences).

### Lentiviral transductions

To produce lentivirus, 900,000 293 FT cells in a 6-well plate were transfected with packaging plasmids (566 ng of pLP1, 266 ng of pLP2, 370 ng of pLP/VSVG) along with 1 µg of lentiviral vector plasmid using 4 µL Lipofectamine 2000 (Thermo Fisher Scientific) as per the manufacturer’s instructions. Medium was refreshed 18 hours later. Virus-containing supernatant was collected 72 hours post transfection, cleared through a 0.45-µm filter, supplemented with 8 µg/ml polybrene (Sigma), and used for infection of target cells. Transductants were selected after one day of recovery. The following antibiotics were used for selection of transductants: puromycin (eHAP 0.4 µg/mL; NCI-H460 1 µg/mL; each for 2-3 days), hygromycin (eHAP 0.4 mg/mL for 3-4 days) and blasticidin (eHAP 8 µg/mL, NCI-H460 2.5 µg/mL; for 5 days).

### Generation of CRISPR full knockout cell lines

Knockouts of *ATRX* and *FAM111A* in eHAP iCas9 and NCI-H460 iCas9 cells were made by transfection of the cell line with synthetic tracrRNA/crRNA using Lipofectamine RNAiMAX (Thermo Fisher Scientific). *ATRX* and *FAM111A* targeting sequences are listed in Supplementary Table 3. Upon transfection cells were incubated with 1 µg/mL doxycycline to induce Cas9 expression for 72 hours, then cells were seeded in 96-well plates to derive as single cell clones. Knockouts of *STN1* and *CTC1* in eHAP iCas9 cells were made by transient transfection of the cell line with lenti-sgRNA-puro (Addgene #104990), targeting sequences are listed in Supplementary Table 3. Upon transfection cells were incubated with 1 µg/mL doxycycline to induce Cas9 expression for 72 hours and selected for 48 hr in puromycin at a concentration of 0.4 µg/mL after 24 hr of transfection. Then cells were seeded in 96-well plates to derive as single cell clones. All stable knockout clones were validated by immunoblotting, sequencing the genomic DNA for the presence of a nonsense/frameshift mutation and screened for iCas9 cutting efficiency (with the BFP/GFP reporter assay).

### Generation of Dox-inducible Cas9 knockout cell lines

Inducible CRISPR knockout cell lines were generated by transducing iCas9 cells with lentivirus produced from the lenti-sgRNA-puro or lenti-sgRNA-hygro constructs (Addgene #104990 and #104991), target sequences of sgRNAs listed in Supplementary Table 3. Inducible knockout of target proteins was confirmed in the pooled cell line by immunoblotting following treatment with 1 µg/mL doxycycline for 96 hr, or where antibodies were unavailable, genomic DNA sequencing of the pooled edited population.

### Generation of endogenous knockin cell lines

High on-target and low off-target scoring sgRNA targeting the exon of interest was designed using Benchling and cloned into the pGL3-U6-sgRNA-PGK-puro plasmid (Addgene #51133). A homology-directed repair (HDR) DNA template containing the mutation of interest surrounded by 250 bp homology arms (Supplementary Table 4) cloned into the pUC19 vector was ordered from Genewiz. 250,000 eHAP iCas9 cells in a 6-well plate were transfected with 4 µg sgRNA plasmid and 2 µg HDR plasmid using Lipofectamine 2000 (Thermo Fisher Scientific). 35 hours post-transfection, cells were selected with 0.4 µg/mL puromycin for 48 hours. 1 µg/mL doxycycline, 1 µM ART558 and 1 µM DNA-PKcs inhibitor were maintained in the media during transfection, selection, and expansion of the cells for a total of 5 days in order to increase HDR rates. Single cell clones were isolated via serial dilution in 96-well plates. HDR was confirmed via Sanger sequencing of the locus of interest. Selected clones were expanded and further validated for protein expression levels using immunoblotting.

### CRISPR/Cas9 screening

Whole genome CRISPR/Cas9 dropout/enrichment screen was performed as described^80^. eHAP iCas9 wild-type or ATRX^KO^ cells were transduced with the lentiviral Brunello library (Addgene #73179-LV, sgRNA only vector). 100 million cells were transduced to achieve a multiplicity of infection (MOI) of 0.4 in three biologically independent transductions, selecting for a coverage of 500 cells per sgRNA. Transduced cells were selected with puromycin (0.4 µg/mL) for 48 hours, after which Cas9 expression was induced with doxycycline (1 µg/mL). Cells were subcultured in doxycycline for the initial 6 days and later subcultured without doxycycline, with representation of 40 million cells (theoretical library coverage of 500 cells per sgRNA) maintained at every step by passaging every two days. Cells were collected both at an early timepoint (day 6 after Dox) and a late timepoint (day 16 after Dox) with the rationale of identifying different sets of genes for which mechanisms of lethality may differ in timings. Samples for sequencing were harvested by washing 60 million cells in PBS, freezing cell pellets, and storing at −80°C.

Genomic DNA was isolated with PureLink Genomic DNA Mini Kit (Thermo Fisher Scientific). Quantity of genomic DNA was measured by Nanodrop and Qubit (Thermo Fisher Scientific). From each sample 200 µg of genomic DNA was then used for library preparation, with one-step amplification of genome-integrated sgRNAs by using P5 mix and P7 barcoded oligonucleotides in a PCR reaction with Ex Taq polymerase (TaKaRa) (Supplementary Table 4). PCR products were purified by agarose gel extraction method using QIAquick Gel Extraction Kit (Qiagen) and additionally purified using MinElute PCR Purification Kit (Qiagen). Concentration of each PCR product was quantified with Qubit dsDNA HS assay (Thermo Fisher Scientific) and 25 ng of PCR product at 4 nM were submitted for sequencing, before which samples went through quality control with Bioanalyzer (Agilent). Libraries were sequenced using HiSeq 4000 with 100 bp reads (30 million reads per sample).

### CRISPR sequencing analysis

Raw data was trimmed by obtaining 20 bp after the first occurrence of ‘‘CACCG’’ in the read sequence. Trimmed reads were then mapped with BWA (version 0.5.9-r16)^81^ to a database of guide sequences for the human CRISPR Brunello lentiviral pooled library downloaded from Addgene (https://www.addgene.org/pooled-library/broadgpp-human-knockout-brunello/) with the parameters ‘‘-l 20 -k 2 -n 2’’. sgRNA counts were obtained after filtering the mapped reads for those that had zero mismatches and mapped to the forward strand of the guide sequence. The MAGeCK ‘test’ command (version 0.5.7)^82^ was used to perform the sgRNA ranking analysis between the relevant conditions with parameters ‘‘–norm-method total–remove-zero both’’. Two pairwise comparisons of samples were done: WT ‘day 6’ vs ATRX-KO ‘day 6’ and WT ‘day 16’ vs ATRX-KO ‘day 16’.

### Gene Ontology and clustering analysis

Dropout hits (top 50 for day 6, top 100 for day 16) were mapped on the protein interaction network through STRING (https://string-db.org) with a minimum required interaction score of 0.4. Subsequently, functional enrichments in both networks were analysed against Gene Ontology (GO) terms Molecular Function, Biological Process and Cellular Component. A whole genome background was assumed for statistical analysis of GO enrichment. Statistics shown (FDR) are p-values corrected for multiple testing within each category using the Benjamin–Hochberg procedure.

### Analysis of sensitivity to DNA replication inhibitors and checkpoint inhibitors

For eHAP iCas9 cells, 200 cells per well were seeded in opaque 96 well plates. For NCI-H460 cells, 400 cells per well were seeded in opaque 96 well plates. Cells were treated the following day with the range of concentrations indicated for each compound. Compounds used are hydroxyurea (Sigma), aphidicolin (Sigma), WEE1 inhibitor MK-1775 (Selleckchem), ATR inhibitor AZD6738 (Selleckchem). After 5 days of treatment, survival was assessed using CellTiter-Glo (Promega) assay, with luminescence measured on a CLARIOStar microplate reader (BMG Labtech). For each cell line luminescence values were normalised against the value of the untreated wells.

### Clonogenic survival assay

400 or 200 eHAP iCas9 cells per well were seeded in 24 well plates (in technical triplicate). Colonies were grown for 6 days. 800 NCI-H460 iCas9 cells per well were seeded in 24 well plates (in technical triplicate). Colonies were grown for 7 days. Colonies were then fixed and stained with 0.5% crystal violet solution with 20% methanol. Plates were scanned and analysed using GelCount (Oxford Optronics). Mean value for eHAP or NCI-H460 iCas9 wild-type cell line with an integrated *NTC* sgRNA or untreated were used for the normalisation.

### IncuCyte proliferation imaging

1000 eHAP iCas9 cells per well were seeded in opaque 96 well plates (in technical triplicate) and grown in an Incucyte S3 System (Sartorius), providing continuous live cell imaging every 3-4 hours up to 5-6 days post-seeding. The “Basic Analyzer” confluence processing analysis tool was used to tailor phase segmentation and quantification. 3-4 images were taken per timepoint per well, and mean value for technical replicates is plotted.

### Two-colour competitive growth assays

eHAP iCas9 ATRX^KO^ or NCI-H460 iCas9 ATRX^KO^ cells were transduced with virus particles expressing sgRNAs either in lenti-sgRNA-GFP-NLS-P2A-puro or lenti-sgRNA-mCherry-NLS-P2A-puro constructs. The sgRNAs used targeted *AAVS1* (control) or a specific gene of interest (Supplementary Table 3). Selected transductants were mixed 1:1 (12,000 cells each) and seeded in a 24-well plate. Cells were imaged for GFP and mCherry signals in an Incucyte S5 System (Sartorius). During the course of the experiment, cells were subcultured when near-confluency was reached. The “Basic Analyzer” fluorescent object ratio analysis tool was used to tailor segmentation and quantification. 9 to 16 images were taken per timepoint per well. Mean values for n=3 biological replicates are plotted.

### DNA fiber assay

eHAP iCas9 cells were pulse labelled with 25 µM CldU and subsequently with 250 µM IdU, for 20 min each. Cells were then harvested by trypsinization. DNA fibre spreads were prepared by spotting 2 mL of cells (500,000 cells/ml in PBS) onto microscope slides followed by lysis with 7 mL of 0.5% SDS, 200 mM Tris-HCl pH 7.4 and 50 mM EDTA. Slides were then tilted to allow a stream of DNA to move slowly toward the bottom of the slide, and DNA spreads were fixed in methanol/acetic acid (3:1). Slides were subsequently denatured in HCl 2.5 M, washed, and blocked in 1% BSA/PBS. HCl-denatured fibre spreads were incubated with rat anti-bromodeoxyuridine (detects CldU, Abcam, ab6326, 1:1,200) and mouse anti-bromodeoxyuridine (detects IdU, B44, BD Biosciences, 1:500) for 1 hour and incubated with anti-rat IgG AlexaFluor 555 and anti-mouse IgG AlexaFluor 488 (both at 1:500, Thermo Fisher Scientific) for 1.5 hours. Slides were mounted in Fluoroshield (Sigma). Images were acquired using a Zeiss Axio Imager M1 microscope, equipped with an ORCA-ER digital camera (Hamamatsu) controlled by Volocity 6.3 software (Perkin Elmer). Fiber length was analysed using ImageJ (NIH). For fork speed analysis, a minimum of 300 fibres were measured per condition in each independent experiment. Fork asymmetry was measured as a percentage of the length ratio of the shortest to the longest fibre of first label origin fibres.

### Whole cell extracts

For whole cell lysates, cells were rinsed with PBS, trypsinised and collected in growth medium. Cells were pelleted by centrifugation at 500 g for 5 min and washed once with PBS. Cell pellets were frozen on dry ice and stored at −80°C. For lysis, cell pellets were thawed on ice and resuspended in RIPA Buffer (10 mM Tris-Cl pH 8.0, 1 mM EDTA, 0.5 mM EGTA, 1% Triton X-100, 0.1% sodium deoxycholate, 0.1% SDS, 140 mM NaCl, 1X phosphatase (PhosSTOP, Roche) and protease (cOmplete, EDTA-free, Roche) inhibitor mixes) and incubated on ice for 20 min. Lysates were sonicated with a probe at medium intensity for 10 seconds in a Soniprep 150 instrument and clarified by centrifugation at 13,000 g for 15 min at 4°C. Protein concentration was determined using the DC Protein Assay (Bio-Rad) according to the manufacturer’s instructions. Proteins were denatured in 2X NuPAGE LDS sample buffer (Invitrogen) and 1% 2-mercaptoethanol (Sigma) for 5 min at 95°C. Lysates were frozen on dry ice and stored at −80°C.

### SDS-PAGE and immunoblotting

Proteins were separated by SDS-PAGE using NuPAGE mini gels (Invitrogen) and transferred onto 0.2 µm pore Nitrocellulose membrane (Amersham Protran; Sigma) using standard procedures. Membranes were blocked with 5% skim milk/TBST (TBS/0.1%Tween-20) for 1 hour at room temperature (RT) and probed with the indicated primary antibodies overnight at 4°C. Membranes were then washed 3 times for 10 min with TBST, incubated with appropriate secondary antibodies conjugated to a horseradish peroxidase (HRP) for 1 hour at RT and washed again 3 times for 10 min with TBST. All incubations were carried out on a horizontal shaker. Immunoblots were developed using Clarity or Clarity Max Western ECL Substrate (Bio-Rad). Chemiluminescence was acquired using a ChemiDoc MP imaging system (Bio-Rad).

### Cell cycle analysis

For EdU/DAPI flow cytometry, eHAP iCas9 cells were labelled for 15 min with 10 µM EdU, fixed in 4% formaldehyde, permeabilized in 0.5% Triton X-100/PBS and washed in 1% BSA/PBS before samples were processed using the Click-iT EdU Flow Cytometry Assay (Thermo Fisher Scientific) with Alexa Fluor 488. DNA was counterstained with DAPI (2 µg/mL). Newly synthesized DNA (EdU) and DNA content (DAPI) were detected using an BD LSRFortessa analyser (Becton Dickinson). Gating of single cells and cell cycle analysis was performed manually using FlowJo.

### Metaphase chromosome spreading

Cells were arrested in metaphase by incubation for 4 hours in medium containing 100 ng/mL nocodazole (Sigma). Mitotic cells were harvested, pelleted and swelled in a hypotonic solution (IMDM:deionized water at 1:3 ratio) for 6 min at RT. Subsequently, cells were fixed dropwise with freshly made Carnoy’s buffer (methanol:glacial acetic acid (3:1)) for 5 min at RT and spun down, this fixation step was repeated three times. The suspension of cells in Carnoy’s buffer was stored at −20°C. For spreading, the cell suspension (100 µL) was dropped on clean glass slides and dried overnight at RT.

### Fluorescence In Situ Hybridization (PNA-FISH)

The chromosome spreads slides were rehydrated in PBS for 5 minutes, fixed in 4% formaldehyde for 5 minutes, treated with 1 mg/ml of pepsin for 10 minutes at 37°C, and fixed in 4% formaldehyde for 5 minutes. Next, slides were dehydrated in 70%, 85%, and 100% (v/v) ethanol for 15 minutes each and then air-dried. Metaphase chromosome spreads were hybridized with a telomeric TelC-Cy5 PNA probe (PNA Bio) and centromeric CENPB-Cy3 probe (PNA Bio) in hybridizing solution (70% formamide, 0.5% blocking reagent (Roche), 10 mM Tris-HCl pH 7.2) for 90 seconds at 80°C followed by 1.5 hours at RT. Slides were washed twice with washing buffer (70% formamide, 10 mM Tris-HCl pH 7.2) for 15 min at RT. Slides were mounted using ProLong Gold antifade with DAPI (Life Technologies). Chromosome images and telomere signals were captured using a Nikon Ti2 microscope fitted with a Prime 95B camera (Photometrics) using Plan Apochromat 100x/1.45 NA Oil objective lens and controlled by Nikon NIS-Elements.

### Immunofluorescence (IF) microscopy coupled to FISH

Cells were grown on #1.5 18 mm glass coverslips. Cells were fixed with 4% formaldehyde in PBS for 15 min at RT. After fixation, cells were washed with PBS three times and then processed for immunofluorescence. Cells were blocked with ADB (Antibody Dilution Buffer; 10% normal goat serum, 0.1% Triton X-100, 0.1% saponin in PBS) for 1 hour at RT. Cells were incubated with primary antibodies (diluted in ADB) for 1 hour at RT, washed three times with PBS and then counterstained with Alexa fluorophore-conjugated secondary antibodies raised in goat (Thermo Fisher Scientific) diluted in ADB, for 1 hour at RT. Cells were then washed three times with PBS. To continue with IF-FISH, cells were fixed again with 4% formaldehyde in PBS for 15 min at RT and then washed twice with PBS. Next, coverslips were dehydrated in 70%, 85%, and 100% (v/v) ethanol for 5 minutes each and then air-dried. Dry coverslips were hybridized with a telomeric TelC-Cy5 PNA probe (PNA Bio) in hybridizing solution (70% formamide, 0.5% blocking reagent (Roche), 10 mM Tris-HCl pH 7.2) for 90 seconds at 80°C followed by 2 hours at RT and washed twice with washing buffer (70% formamide, 10 mM Tris-HCl pH 7.2) for 15 min at RT. The coverslips were mounted onto glass slides with ProLong Gold antifade with DAPI (Life Technologies). Images were acquired using a Nikon Ti2 microscope fitted with a CSU-W1 spinning disk confocal unit (Yokogawa) and a Prime 95B camera (Photometrics) using Nikon CFI Apochromat LWD Lambda S 40x/1.15 WI objective lens and controlled by Nikon NIS-Elements. Following acquisition, images were imported into Fiji (NIH) for automated foci counting and colocalization quantitation.

### C-circle assay

The C-circle assay protocol was adapted from Henson et al^83^, by using the quick C-circle preparation (QCP) protocol. DNA concentration was measured by fluorimetry using the Qubit dsDNA HS Assay (Thermo Fisher Scientific). Samples were pre-diluted in QCP lysis buffer at 30 ng/µl. 30 ng of DNA were diluted to 10 µl in 10 mM Tris-HCl pH 7.6 and mixed with 9.25 µl of Rolling Circle Master Mix (RCMM) (8.65 mM DTT, 2.16X phi29 buffer, 8.65 µg/mL BSA, 0.216% Tween-20 and 2.16 mM of each dATP, dCTP, dGTP and dTTP) and 0.75 µl of phi29 DNA Polymerase (Thermo Fisher Scientific). Rolling circle amplification was performed by incubating samples in a thermocycler at 30°C for 8 hours, polymerase was inactivated at 70°C for 20 min and then kept at 8-10°C. For slot blot detection, samples were blotted onto Amersham Hybond N+ positively charged nylon membranes (GE Healthcare) under native conditions. After crosslinking, membranes were hybridized with non-radioactive 3’ DIG labelled probes, as previously described^84^. Membranes were developed using anti-DIG-AP (Roche) and CDP-Star substrate (Roche). Chemiluminescence was acquired using a ChemiDoc MP imaging system (Bio-Rad). Membranes were stripped and re-hybridised with 3’DIG labelled Alu probe as a loading control.

### Native FISH (ssCCCTAA/ssTTAGGG)

Cells were grown on coverslips in 12-well plates or 96-well microplates (PerkinElmer) and fixed in 2% formaldehyde for 15 min at RT. After fixation, cells were washed with PBS twice and then processed for native FISH. Cells were incubated with 250 μg/mL RNaseA in ADB blocking solution (Antibody Dilution Buffer; 10% normal goat serum, 0.1% Triton X-100, 0.1% saponin in PBS) for 1 h at 37°C. Next, coverslips/plates were dehydrated in 70%, 85%, and 100% (v/v) ethanol for 5 minutes each and then air-dried. Dry coverslips/plates were hybridized with a telomeric TelG-TAMRA PNA probe (PNA Bio) (for ssCCCTAA) or TelC-Cy5 PNA probe (PNA Bio) (for ssTTAGGG) in hybridizing solution (70% formamide, 1 mg/mL blocking reagent (Roche), 10 mM Tris-HCl pH 7.5) for 2 hours at RT and washed twice with PBS. Nuclei were stained with incubation for 5 min with DAPI in PBS, followed by three washes with PBS. Coverslips were mounted in ProLong Gold antifade with DAPI (Life Technologies) and imaged in a Nikon Ti2 microscope fitted with a Prime 95B camera (Photometrics) using Plan Apochromat 60x/1.4 NA Oil objective lens and controlled by Nikon NIS-Elements. Alternatively, plates were sealed and scanned using an Opera Phenix Plus High-Content Screening System (PerkinElmer), where images were captured using 40x objective and analysed using Harmony High-Content Imaging and Analysis Software (PerkinElmer).

### Telomere G-overhang assay

For the analysis of human telomeric DNA by in gel hybridization, the protocol was adapted from Wu et al^15^. Cells were trypsinised and resuspended in PBS. They were then mixed 1:1 with 2% agarose (Lonza) in PBS to obtain a concentration of 1 million cells per plug. Plugs were digested with 1 mg/mL Proteinase K in digest buffer (100 mM EDTA pH 8.0, 0.2% sodium deoxycholate, 1% sarcosyl) overnight at 50°C. Plugs were then washed four times for 1 hr each in TE buffer, with 1 mM PMSF in the last wash, once with sterile water for 30 min, and with restriction enzyme buffer (CutSmart) for 30 min. Plugs were incubated with 60 units MboI in restriction enzyme buffer overnight at 37°C. The following day, the plugs were rinsed once in TE, and once in 0.5X TBE, and loaded onto a 1% agarose/0.5X TBE gel and separated by PFGE for 18–24 hr (CHEF DR III, Bio-Rad) in 0.5X TBE. The gels were dried and prehybridized in Church mix for 1 hr at 50°C. Hybridization with radioactively labelled γ-32P-ATP [AACCCT]_4_ was performed overnight at 50°C in Church mix. The gel was washed at 55°C three times for 30 min each in 4X SSC and once for 30 min in 0.1% SDS/4X SSC. The gel was exposed to a phosphorimaging plate (GE Healthcare) and scanned using Typhoon FLA 9500 (GE Healthcare). The gel was subsequently denatured in 1.5 M NaCl, 0.5 M NaOH for 30 min, neutralised with two 15 min washes in 0.5 M Tris-HCl pH7.5, 3M NaCl, and pre-hybridised with the same probe overnight at 55°C. The gel was washed, exposed, and imaged as above. The single-stranded overhang signal in the native gel relative to the total telomeric DNA in the denatured gel was quantified using Fiji (NIH).

### SILAC Labeling

eHAP cells were grown in IMDM for SILAC (Thermo Fisher Scientific) supplemented with 10% (v/v) SILAC dialyzed FBS (Gemini Biosciences) and 1% penicillin/streptomycin. Cells designated as “heavy” were cultured in media supplemented with L-Lysine (13C6/15N2), L-Arginine (13C6/15N4) (both from Cambridge Isotope Laboratories) and L-Proline, while cells designated as “light” were cultured in media supplemented with unlabelled L-Lysine, L-Arginine and L-Proline. Cells were passaged for at least 10-12 population doublings, where a labelling efficiency test was performed to confirm label incorporation.

### iPOND

Asynchronous eHAP WT and ATRX^KO^ cultures were pulsed with 10 uM EdU for 10 min. For HU chase, cells were pulsed with 10 uM EdU for 10 min, washed once in pre-equilibrated chase medium, and treated with 0.5 mM HU for 60 min. For thymidine chase, cells were pulsed with 10 uM EdU for 10 min, washed once in pre-equilibrated chase medium, and treated with 10 uM thymidine for 60 min. Cells were immediately crosslinked with 1% formaldehyde in PBS for 20 min at RT, quenched for 20 min with 0.125 M glycine in PBS, collected by scraping, and washed three times in cold PBS. For SILAC, heavy and light cells were pooled 1:1 at this stage. Pellets were resuspended in 10 mL permeabilization buffer (0.25% (v/v) Triton X-100 in PBS), incubated at RT for 30 min, and spun at 1000g for 5 min. The pellets were washed in 10 mL 0.5% (w/v) BSA in PBS, spun at 1000g for 5 min and resuspended in Click Reaction cocktail (in a total volume of 5 mL in PBS: 10 uM biotin azide, 10 mM sodium ascorbate, 2 mM CuSO4), incubated for 2 hours at RT. The pellets were then washed in 0.5% (w/v) BSA in PBS, spun at 1000g for 5 min, resuspended in 700 uL Nuclear Lysis buffer (10 mM Tris-Cl pH 8.0, 1 mM EDTA, 0.5 mM EGTA, 1% Triton X-100, 0.1% sodium deoxycholate, 0.1% SDS, 140 mM NaCl, 1X phosphatase (PhosSTOP, Roche) and protease (cOmplete, EDTA-free, Roche) inhibitor mixes), incubated on ice for 15 min, sonicated in a Bioruptor Pico instrument (Diagenode) for a total of 25 cycles (30’’on/off) and spun in a microcentrifuge at maximum speed for 10 min. Lysate was diluted 1:1 with cold PBS and subjected to pull-down with 50 uL Sera-Mag SpeedBeads Neutravidin-Coated Magnetic Particles (Cytiva) overnight at 4°C. Beads were washed four times with Lysis buffer, eluted for 20 min at 95°C in 2X NuPAGE LDS sample buffer (Invitrogen) and 2% 2-mercaptoethanol (Sigma), and eluate was run 3 cm into a 10% NuPAGE Bis-Tris gel (Invitrogen) for mass spectrometry analysis.

### Immunoprecipitation

eHAP ATRX KO cells stably complemented with ATRX-HA were lysed by incubating them for 40 min under constant agitation in IP buffer (10 mM Tris pH 7.5, 150 mM NaCl, 0.5 mM EDTA, 2.5 mM MgCl2, 0.5% Triton X-100) complemented with 1X phosphatase (PhosSTOP, Roche) and protease (cOmplete, EDTA-free, Roche) inhibitor mixes and benzonase (E1014, Sigma). Cells were then centrifuged at 15,000 x g for 15 min and the cleared cell lysates were diluted in dilution buffer (10 mM Tris pH 7.5, 150 mM NaCl, 0.5 mM EDTA, 2.5 mM MgCl2) complemented with 1X phosphatase (PhosSTOP, Roche) and protease (cOmplete, EDTA-free, Roche) inhibitor mixes. Dilluted lysates were immunoprecipitated using 30 μL of Pierce anti-HA magnetic beads (88826, Thermo) overnight at 4°C. Beads were washed three times for 10 minutes with dilution buffer buffer containing 0.05% NP-40 and eluted and boiled in 2X NuPAGE LDS sample buffer (Invitrogen) and 1% 2-mercaptoethanol (Sigma).

### Chromatin fractionation

eHAP iCas9 cells were seeded 36-48 hr prior to collection. Asynchronous cultures were trypsinised and resuspended in ice-cold PBS. 3 million cells were kept on ice for whole cell lysate control. 3 million cells were spun down for 5 min at 500 g, and resuspended in 200 µL CSK buffer (10 mM PIPES pH 7.0, 100 mM NaCl, 300 mM sucrose, 1.5 mM MgCl2, 5 mM EDTA, 0.5% Triton X-100, 1X phosphatase (PhosSTOP, Roche) and protease (cOmplete, EDTA-free, Roche) inhibitor mixes) and incubated on ice for 10 min. Cells were spun down at full speed for 10 seconds and the supernatant (soluble fraction) was collected. The chromatin pellet was washed in 500 µL of CSK buffer. Chromatin pellets were resuspended in 200 µL 1X NuPAGE LDS sample buffer (Invitrogen) and 1% 2-mercaptoethanol (Sigma). 50 µL of 4X NuPAGE LDS sample buffer (Invitrogen) and 4% 2-mercaptoethanol (Sigma) was added to 150 µL whole cell lysate samples. All samples were boiled at 95°C for 10 min and sonicated with a probe at medium intensity for 10 seconds in a Soniprep 150 instrument. 20 µL of each fraction was loaded and subjected to SDS-PAGE as above.

### Statistical analysis

Sample number (n) indicates the number of independent biological samples in each experiment and are indicated in figure legends. Statistical details of each experiment (including the statistical tests used, exact value of n) can be found in the figure legends. Prism 9/10 (GraphPad) and RStudio were used for statistical analysis: one-way ANOVA, followed by Tukey’s multiple comparison test was used unless stated otherwise (P < 0.05 (*), P < 0.01 (**), P < 0.001 (***), P < 0.0001 (****)).

## Data availability

Further information and requests for resources and reagents should be directed to and will be fulfilled by the lead contact, Simon J. Boulton. Any additional information required to reanalyse the data reported in this paper is available upon request.

## References

1. Saxena, S. & Zou, L. Hallmarks of DNA replication stress. Mol Cell 82, 2298–2314 (2022).

2. Zou, L. & Elledge, S.J. Sensing DNA damage through ATRIP recognition of RPA-ssDNA complexes. Science 300, 1542–8 (2003).

3. Cortez, D., Guntuku, S., Qin, J. & Elledge, S.J. ATR and ATRIP: partners in checkpoint signaling. Science 294, 1713–6 (2001).

4. Delacroix, S., Wagner, J.M., Kobayashi, M., Yamamoto, K. & Karnitz, L.M. The Rad9-Hus1-Rad1 (9-1-1) clamp activates checkpoint signaling via TopBP1. Genes Dev 21, 1472–7 (2007).

5. Haahr, P. et al. Activation of the ATR kinase by the RPA-binding protein ETAA1. Nat Cell Biol 18, 1196–1207 (2016).

6. Bass, T.E. et al. ETAA1 acts at stalled replication forks to maintain genome integrity. Nat Cell Biol 18, 1185–1195 (2016).

7. Matsuoka, S. et al. ATM and ATR substrate analysis reveals extensive protein networks responsive to DNA damage. Science 316, 1160–6 (2007).

8. Liu, Q. et al. Chk1 is an essential kinase that is regulated by Atr and required for the G(2)/M DNA damage checkpoint. Genes Dev 14, 1448–59 (2000).

9. Saldivar, J.C., Cortez, D. & Cimprich, K.A. The essential kinase ATR: ensuring faithful duplication of a challenging genome. Nat Rev Mol Cell Biol 18, 622–636 (2017).

10. Sfeir, A. et al. Mammalian telomeres resemble fragile sites and require TRF1 for efficient replication. Cell 138, 90–103 (2009).

11. Hom, R.A. & Wuttke, D.S. Human CST Prefers G-Rich but Not Necessarily Telomeric Sequences. Biochemistry 56, 4210–4218 (2017).

12. Chen, L.Y., Redon, S. & Lingner, J. The human CST complex is a terminator of telomerase activity. Nature 488, 540–4 (2012).

13. Zaug, A.J. et al. CST does not evict elongating telomerase but prevents initiation by ssDNA binding. Nucleic Acids Res 49, 11653–11665 (2021).

14. Wang, F. et al. Human CST has independent functions during telomere duplex replication and C-strand fill-in. Cell Rep 2, 1096–103 (2012).

15. Wu, P., Takai, H. & de Lange, T. Telomeric 3’ overhangs derive from resection by Exo1 and Apollo and fill-in by POT1b-associated CST. Cell 150, 39–52 (2012).

16. Takai, H., Aria, V., Borges, P., Yeeles, J.T.P. & de Lange, T. CST-polymerase alpha-primase solves a second telomere end-replication problem. Nature 627, 664–670 (2024).

17. Stewart, J.A. et al. Human CST promotes telomere duplex replication and general replication restart after fork stalling. EMBO J 31, 3537–49 (2012).

18. Chastain, M. et al. Human CST Facilitates Genome-wide RAD51 Recruitment to GC-Rich Repetitive Sequences in Response to Replication Stress. Cell Rep 16, 2048 (2016).

19. Wang, Y. & Chai, W. Pathogenic CTC1 mutations cause global genome instabilities under replication stress. Nucleic Acids Res 46, 3981–3992 (2018).

20. Lei, K.H. et al. Crosstalk between CST and RPA regulates RAD51 activity during replication stress. Nat Commun 12, 6412 (2021).

21. Lyu, X. et al. Human CST complex protects stalled replication forks by directly blocking MRE11 degradation of nascent-strand DNA. EMBO J 40, e103654 (2021).

22. Zhang, M. et al. Mammalian CST averts replication failure by preventing G-quadruplex accumulation. Nucleic Acids Res 47, 5243–5259 (2019).

23. Mirman, Z. et al. 53BP1-RIF1-shieldin counteracts DSB resection through CST-and Polalpha-dependent fill-in. Nature 560, 112–116 (2018).

24. Gibbons, R.J., Picketts, D.J., Villard, L. & Higgs, D.R. Mutations in a putative global transcriptional regulator cause X-linked mental retardation with alpha-thalassemia (ATR-X syndrome). Cell 80, 837–45 (1995).

25. Cerami, E. et al. The cBio cancer genomics portal: an open platform for exploring multidimensional cancer genomics data. Cancer Discov 2, 401–4 (2012).

26. Gao, J. et al. Integrative analysis of complex cancer genomics and clinical profiles using the cBioPortal. Sci Signal 6, pl1 (2013).

27. Goldberg, A.D. et al. Distinct factors control histone variant H3.3 localization at specific genomic regions. Cell 140, 678–91 (2010).

28. Lewis, P.W., Elsaesser, S.J., Noh, K.M., Stadler, S.C. & Allis, C.D. Daxx is an H3.3-specific histone chaperone and cooperates with ATRX in replication-independent chromatin assembly at telomeres. Proc Natl Acad Sci U S A 107, 14075–80 (2010).

29. Drane, P., Ouararhni, K., Depaux, A., Shuaib, M. & Hamiche, A. The death-associated protein DAXX is a novel histone chaperone involved in the replication-independent deposition of H3.3. Genes Dev 24, 1253–65 (2010).

30. Law, M.J. et al. ATR-X syndrome protein targets tandem repeats and influences allele-specific expression in a size-dependent manner. Cell 143, 367–78 (2010).

31. Wong, L.H. et al. ATRX interacts with H3.3 in maintaining telomere structural integrity in pluripotent embryonic stem cells. Genome Res 20, 351–60 (2010).

32. Voon, H.P. et al. ATRX Plays a Key Role in Maintaining Silencing at Interstitial Heterochromatic Loci and Imprinted Genes. Cell Rep 11, 405–18 (2015).

33. Sadic, D. et al. Atrx promotes heterochromatin formation at retrotransposons. EMBO Rep 16, 836–50 (2015).

34. Teng, Y.C. et al. ATRX promotes heterochromatin formation to protect cells from G-quadruplex DNA-mediated stress. Nat Commun 12, 3887 (2021).

35. Truch, J. et al. The chromatin remodeller ATRX facilitates diverse nuclear processes, in a stochastic manner, in both heterochromatin and euchromatin. Nat Commun 13, 3485 (2022).

36. Clynes, D. et al. ATRX dysfunction induces replication defects in primary mouse cells. PLoS One 9, e92915 (2014).

37. Huh, M.S. et al. Stalled replication forks within heterochromatin require ATRX for protection. Cell Death Dis 7, e2220 (2016).

38. Nguyen, D.T. et al. The chromatin remodelling factor ATRX suppresses R-loops in transcribed telomeric repeats. EMBO Rep 18, 914–928 (2017).

39. Wang, Y. et al. G-quadruplex DNA drives genomic instability and represents a targetable molecular abnormality in ATRX-deficient malignant glioma. Nat Commun 10, 943 (2019).

40. Juhasz, S., Elbakry, A., Mathes, A. & Lobrich, M. ATRX Promotes DNA Repair Synthesis and Sister Chromatid Exchange during Homologous Recombination. Mol Cell 71, 11–24 e7 (2018).

41. Cotta-Ramusino, C. et al. A DNA damage response screen identifies RHINO, a 9-1-1 and TopBP1 interacting protein required for ATR signaling. Science 332, 1313–7 (2011).

42. Drosopoulos, W.C., Kosiyatrakul, S.T., Yan, Z., Calderano, S.G. & Schildkraut, C.L. Human telomeres replicate using chromosome-specific, rather than universal, replication programs. J Cell Biol 197, 253–66 (2012).

43. Loe, T.K. et al. Telomere length heterogeneity in ALT cells is maintained by PML-dependent localization of the BTR complex to telomeres. Genes Dev 34, 650–662 (2020).

44. Mazzucco, G. et al. Telomere damage induces internal loops that generate telomeric circles. Nat Commun 11, 5297 (2020).

45. Huang, C., Jia, P., Chastain, M., Shiva, O. & Chai, W. The human CTC1/STN1/TEN1 complex regulates telomere maintenance in ALT cancer cells. Exp Cell Res 355, 95–104 (2017).

46. Lee, J. et al. Extrachromosomal telomere DNA derived from excessive strand displacements. Proc Natl Acad Sci U S A 121, e2318438121 (2024).

47. Lovejoy, C.A., Takai, K., Huh, M.S., Picketts, D.J. & de Lange, T. ATRX affects the repair of telomeric DSBs by promoting cohesion and a DAXX-dependent activity. PLoS Biol 18, e3000594 (2020).

48. Zhong, S. et al. A role for PML and the nuclear body in genomic stability. Oncogene 18, 7941–7 (1999).

49. Carbone, R., Pearson, M., Minucci, S. & Pelicci, P.G. PML NBs associate with the hMre11 complex and p53 at sites of irradiation induced DNA damage. Oncogene 21, 1633–40 (2002).

50. Marchesini, M. et al. PML is required for telomere stability in non-neoplastic human cells. Oncogene 35, 1811–21 (2016).

51. Jiang, H. et al. BLM helicase unwinds lagging strand substrates to assemble the ALT telomere damage response. Mol Cell 84, 1684–1698 e9 (2024).

52. Buisson, R., Boisvert, J.L., Benes, C.H. & Zou, L. Distinct but Concerted Roles of ATR, DNA-PK, and Chk1 in Countering Replication Stress during S Phase. Mol Cell 59, 1011–24 (2015).

53. Kim, S.T., Lim, D.S., Canman, C.E. & Kastan, M.B. Substrate specificities and identification of putative substrates of ATM kinase family members. J Biol Chem 274, 37538–43 (1999).

54. Liang, J. et al. Genome-Wide CRISPR-Cas9 Screen Reveals Selective Vulnerability of ATRX-Mutant Cancers to WEE1 Inhibition. Cancer Res 80, 510–523 (2020).

55. Sirbu, B.M. et al. Identification of proteins at active, stalled, and collapsed replication forks using isolation of proteins on nascent DNA (iPOND) coupled with mass spectrometry. J Biol Chem 288, 31458–67 (2013).

56. Dungrawala, H. et al. The Replication Checkpoint Prevents Two Types of Fork Collapse without Regulating Replisome Stability. Mol Cell 59, 998–1010 (2015).

57. Wessel, S.R., Mohni, K.N., Luzwick, J.W., Dungrawala, H. & Cortez, D. Functional Analysis of the Replication Fork Proteome Identifies BET Proteins as PCNA Regulators. Cell Rep 28, 3497–3509 e4 (2019).

58. Mukherjee, C. et al. RIF1 promotes replication fork protection and efficient restart to maintain genome stability. Nat Commun 10, 3287 (2019).

59. Alabert, C. et al. Nascent chromatin capture proteomics determines chromatin dynamics during DNA replication and identifies unknown fork components. Nat Cell Biol 16, 281–93 (2014).

60. Unger, S. et al. FAM111A mutations result in hypoparathyroidism and impaired skeletal development. Am J Hum Genet 92, 990–5 (2013).

61. Kojima, Y. et al. FAM111A protects replication forks from protein obstacles via its trypsin-like domain. Nat Commun 11, 1318 (2020).

62. Hoffmann, S. et al. FAM111 protease activity undermines cellular fitness and is amplified by gain-of-function mutations in human disease. EMBO Rep 21, e50662 (2020).

63. Rios-Szwed, D.O. et al. FAM111A regulates replication origin activation and cell fitness. Life Sci Alliance 6(2023).

64. Eustermann, S. et al. Combinatorial readout of histone H3 modifications specifies localization of ATRX to heterochromatin. Nat Struct Mol Biol 18, 777–82 (2011).

65. Iwase, S. et al. ATRX ADD domain links an atypical histone methylation recognition mechanism to human mental-retardation syndrome. Nat Struct Mol Biol 18, 769–76 (2011).

66. Hoelper, D., Huang, H., Jain, A.Y., Patel, D.J. & Lewis, P.W. Structural and mechanistic insights into ATRX-dependent and -independent functions of the histone chaperone DAXX. Nat Commun 8, 1193 (2017).

67. Xue, Y. et al. The ATRX syndrome protein forms a chromatin-remodeling complex with Daxx and localizes in promyelocytic leukemia nuclear bodies. Proc Natl Acad Sci U S A 100, 10635–40 (2003).

68. Tang, J. et al. A novel transcription regulatory complex containing death domain-associated protein and the ATR-X syndrome protein. J Biol Chem 279, 20369–77 (2004).

69. Wasylishen, A.R. et al. Daxx maintains endogenous retroviral silencing and restricts cellular plasticity in vivo. Sci Adv 6, eaba8415 (2020).

70. Pinto, L.M. et al. DAXX promotes centromeric stability independently of ATRX by preventing the accumulation of R-loop-induced DNA double-stranded breaks. Nucleic Acids Res 52, 1136–1155 (2024).

71. Tomaska, L., Cesare, A.J., AlTurki, T.M. & Griffith, J.D. Twenty years of t-loops: A case study for the importance of collaboration in molecular biology. DNA Repair (Amst) 94, 102901 (2020).

72. Surovtseva, Y.V. et al. Conserved telomere maintenance component 1 interacts with STN1 and maintains chromosome ends in higher eukaryotes. Mol Cell 36, 207–18 (2009).

73. Miyake, Y. et al. RPA-like mammalian Ctc1-Stn1-Ten1 complex binds to single-stranded DNA and protects telomeres independently of the Pot1 pathway. Mol Cell 36, 193–206 (2009).

74. Huang, C., Dai, X. & Chai, W. Human Stn1 protects telomere integrity by promoting efficient lagging-strand synthesis at telomeres and mediating C-strand fill-in. Cell Res 22, 1681–95 (2012).

75. van Schendel, R., Romeijn, R., Buijs, H. & Tijsterman, M. Preservation of lagging strand integrity at sites of stalled replication by Pol alpha-primase and 9-1-1 complex. Sci Adv 7(2021).

76. Panda, D., Fernandez, D.J., Lal, M., Buehler, E. & Moss, B. Triad of human cellular proteins, IRF2, FAM111A, and RFC3, restrict replication of orthopoxvirus SPI-1 host-range mutants. Proc Natl Acad Sci U S A 114, 3720–3725 (2017).

77. Nie, M. et al. FAM111A induces nuclear dysfunction in disease and viral restriction. EMBO Rep 22, e50803 (2021).

78. Palani, S. et al. Dimerization-dependent serine protease activity of FAM111A prevents replication fork stalling at topoisomerase 1 cleavage complexes. Nat Commun 15, 2064 (2024).

79. Hewitt, G. et al. Defective ALC1 nucleosome remodeling confers PARPi sensitization and synthetic lethality with HRD. Mol Cell 81, 767–783 e11 (2021).

80. Doench, J.G. et al. Optimized sgRNA design to maximize activity and minimize off-target effects of CRISPR-Cas9. Nat Biotechnol 34, 184–191 (2016).

81. Li, H. et al. The Sequence Alignment/Map format and SAMtools. Bioinformatics 25, 2078–9 (2009).

82. Li, W. et al. MAGeCK enables robust identification of essential genes from genome-scale CRISPR/Cas9 knockout screens. Genome Biol 15, 554 (2014).

83. Henson, J.D. et al. The C-Circle Assay for alternative-lengthening-of-telomeres activity. Methods 114, 74–84 (2017).

84. Idilli, A.I., Segura-Bayona, S., Lippert, T.P. & Boulton, S.J. A C-circle assay for detection of alternative lengthening of telomere activity in FFPE tissue. STAR Protoc 2, 100569 (2021).

